# Embryological cellular origins and hypoxia-mediated mechanisms in *PIK3CA*-Driven refractory vascular malformations

**DOI:** 10.1101/2024.10.16.618777

**Authors:** Sota Torii, Keiki Nagaharu, Nanako Nakanishi, Hidehito Usui, Yumiko Hori, Katsutoshi Hirose, Satoru Toyosawa, Eiichi Morii, Mitsunaga Narushima, Yoshiaki Kubota, Osamu Nakagawa, Kyoko Imanaka-Yoshida, Kazuaki Maruyama

## Abstract

Congenital vascular malformations, affecting 0.5% of the population, often occur in the head and neck, complicating treatment due to the critical functions in these regions. Our previous research identified distinct developmental origins for blood and lymphatic vessels in these areas, tracing them to the cardiopharyngeal mesoderm (CPM), which contributes to the development of the head, neck, and cardiovascular system in both mouse and human embryos.

In this study, we investigated the pathogenesis of these malformations by expressing Pik3ca^H1047R^ in the CPM. Mice expressing Pik3ca^H1047R^ in the CPM developed vascular abnormalities restricted to the head and neck. Single-cell RNA sequencing revealed that Pik3ca^H1047R^ upregulates *Vegf-a* expression in endothelial cells through HIF-mediated hypoxia signaling. Human samples supported these findings, showing elevated HIF-1α and VEGF-A in malformed vessels. Notably, inhibition of HIF-1α and VEGF-A in the mouse model significantly reduced abnormal vasculature. These results highlight the role of embryonic origins and hypoxia-driven mechanisms in vascular malformations, providing a foundation for the development of therapies targeting these difficult-to-treat conditions.

## Introduction

Vascular malformations, including lymphatic (LMs) and venous malformations (VMs), are chronic, often debilitating conditions that arise during embryonic development^1–3^. These malformations commonly present with significant clinical symptoms, especially in the head and neck, where approximately 80% of LMs, 90% of capillary malformations, and 40% of VMs occur ^4,5^. In these cases, they can impair vital functions such as breathing and feeding, as well as affect appearance. Vascular malformations in the head and neck region are particularly challenging to treat and have been classified as intractable diseases, setting them apart from vascular malformations occurring in other parts of the body^6–8^. Standard treatments, including surgical excision and sclerotherapy, are often employed but not always effective. Thus, there is an urgent need for novel therapies that target the underlying disease mechanisms. Vascular malformations are generally categorized into low-flow (venous, lymphatic, and capillary) and fast-flow (arteriovenous) lesions, with low-flow malformations being the most common^8,9^. *PIK3CA* mutations are found in approximately 85% of LMs^4,10–12^ and 20% of VMs^13–16^, though this varies between studies. PIK3CA encodes the p110α lipid kinase, responsible for converting PIP2 to PIP3, which activates key downstream signaling pathways. This kinase is crucial for the development of blood and lymphatic vessels^17–19^, and is activated by receptor tyrosine kinases, including vascular endothelial growth factor (VEGF) receptors and the TIE2/TEK receptor. Studies in mice have shown that p110α is essential for normal blood and lymphatic vessel development. Conversely, expression of the *PIK3CA^H1047R^* mutation in endothelial cells (ECs) has been shown to cause vascular malformations^12,20^.

VMs and LMs are frequently driven by mutations in the helical (E542K, E545K) and kinase (H1047R, H1047L) domains of p110α, identical to those observed in cancer and genetic syndromes marked by tissue overgrowth, such as *PIK3CA*-related overgrowth spectrum (PROS)^21,22^. These somatic *PIK3CA* mutations activate key downstream pathways, most notably the AKT-mTOR pathway, which is thought to play a significant role in disease progression. However, the exact mechanisms remain unclear, with additional pathways, such as ERK and transforming growth factor-α (TGF-α), also being implicated^23–27^. The effects of these mutations may not be limited to ECs because abnormal ECs can interact with surrounding tissues, potentially driving fibrosis^24,25^. While mTOR inhibitors like rapamycin and its analogs (sirolimus, everolimus) have been effective in reducing the symptoms of vascular malformations, complete lesion regression is rare. This suggests the involvement of additional pathways beyond mTOR, underscoring the need for more potent therapies^20,28–31^.

Our research highlights the unique pathophysiology of vascular malformations, particularly their development in the head and neck, where ECs arise from the *Islet1* (*Isl1*)^+^ cardiopharyngeal mesoderm (CPM)^32–35^.

Herein, we demonstrated that the expression of Pik3ca^H1047R^ in the CPM generates a model mouse that closely mimics human pathologies. Notably, although the CPM differentiates into a broad range of cell types, the effects of Pik3ca^H1047R^ were specifically confined to veins and lymphatic vessels. Single-cell RNA sequencing (scRNA-seq) of FACS-sorted eYFP^+^ cells from *Isl1-Cre; Pik3ca^H1047R^; R26R-eYFP* embryos and Bulk RNA-seq of PIK3CA^H1047R^- expressing human umbilical vein ECs (HUVECs) and patient-derived ECs revealed hypoxia-driven metabolic shifts and enhanced angiogenesis, including elevated Vegf-a production, contributing to vascular malformation pathogenesis. Immunohistochemical analysis of human vascular malformation samples confirmed the elevated HIF-1α and VEGF-A levels in abnormal vessels, reinforcing this hypothesis. Additionally, inhibiting HIF-1α and VEGF-A signaling in our mouse model significantly suppressed lesion formation, suggesting a potential therapeutic strategy for these intractable conditions.

This study offers new insights into the anatomical and molecular mechanisms underlying vascular malformations, highlighting the significance of developmental origins and hypoxia-mediated signaling pathways. These findings provide a promising foundation for developing novel therapeutic strategies aimed at targeting these complex and currently untreatable vascular anomalies.

## Results

### Expression of Pik3ca^H1047R^ in endothelial cells recapitulates human low-flow vascular malformations

To assess whether Pik3ca^H1047R^ expression can recapitulate the VMs and LMs observed in humans, we utilized a transgenic mouse model, LoxP-STOP-LoxP–Pik3ca^H1047R^ (*R26R- Pik3ca^H1047R^*). This model enables tissue-specific activation of Pik3ca^H1047R^ through Cre-loxP technology. By inducing Pik3ca^H1047R^ during embryonic development using various Cre-expressing strains, we observed that crossing with *Tie2-Cre*, which drives Cre expression in ECs, resulted in widespread dilated, blood-filled vasculature by embryonic day (E)13.5 **(Figure 1A)**. Sagittal section co-immunostaining at E13.5 with EC markers, namely platelet endothelial cell adhesion molecule (PECAM) and VEGF receptor 3 (VEGFR3), as well as the lymphatic marker Prospero homeobox protein 1 (Prox1), revealed a dilated cardinal vein surrounded by an increased number of blood-filled PECAM^+^/Prox1^+^/VEGFR3^+^ lymphatic vessels in *Tie2-Cre; R26R-Pik3ca^H1047R^* mutant embryos **(Figure 1B-F)**. Although less prominent than in the cardinal vein, dilated PECAM^+^/Prox1^-^/partially VEGFR3^+^ blood vessels were also observed extending from the pharyngeal arches to the mandible in mutant embryos (**Figure 1B, G-I**). The overall number of PECAM^+^ vessels did not significantly differ between control and mutant embryos (**Figure 1J**). In the liver, we observed an increase in spongy, dilated PECAM^+^/Prox1^-^/partially VEGFR3^+^ veins connected to the inferior vena cava (**Figure 1B, K-N**). Additionally, in the brain, there was an increase in tortuous and slightly dilated PECAM^+^/Prox1^-^/VEGFR3^+^ blood vessels (**Figure 1B, O-R**).

**Figure 1.**
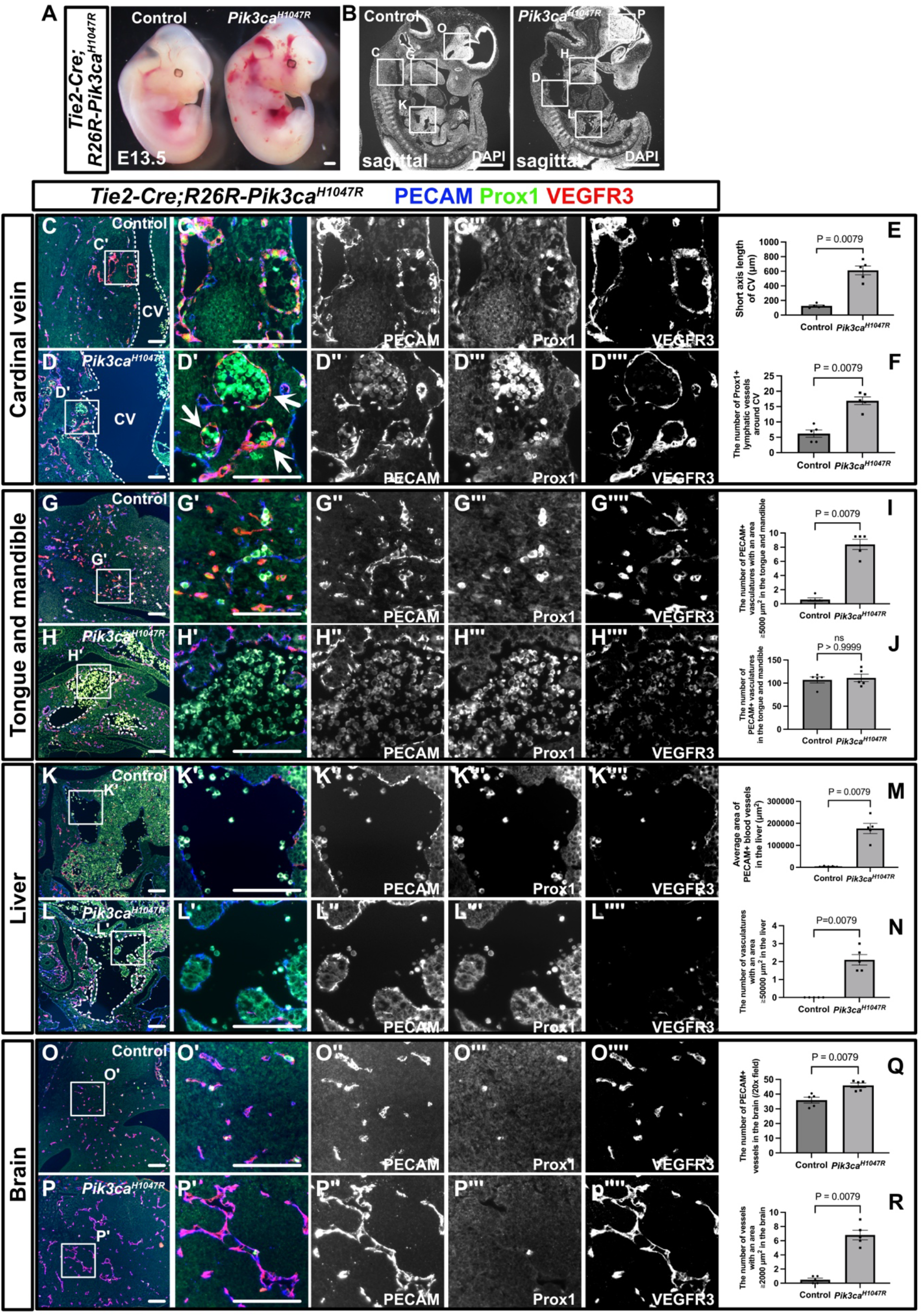
*Tie2-Cre; R26R-Pik3ca^H1047R^* embryos mimic human venous and lymphatic malformations. **(A)** Gross morphology of control and *Tie2-Cre; R26R-Pik3ca^H1047R^* mutant embryos at E13.5. **(B, C-D’’”, G-H’’”, K-L’’”, O-P’’”)** Immunostaining of sagittal sections with the indicated antibodies. **(B-F)** In the mutants, the cardinal vein (CV) is dilated **(E)**, and there is an increase in blood-filled PRECAM^+^/ Prox1^+^/VEGFR3^+^ lymphatic vessels surrounding the CV (white arrows) **(D)**. **(B, G-J)** In the mandible-tongue region, there is an expansion of blood-filled PECAM^+^/Prox1^-^/partially VEGFR3^+^ blood vessels with an area greater than 5000 μm^2^ (white dotted lines) **(H)**. No significant increase in the total number of PECAM^+^ vessels was observed **(J)**. **(B, K-N)** In the liver, irregularly shaped, dilated PECAM^+^/Prox1^-^/partially VEGFR3^+^ blood vessels with an area greater than 5000 μm^2^ (white dotted lines) were observed **(M, N)**. **(O, P)** Similarly, in the brain, irregularly shaped and dilated PECAM^+^/Prox1^-^/VEGFR3^+^ blood vessels with an area greater than 2000 μm^2^ were observed **(Q, R)**. Each dot represents a value obtained from one sample. Scale bars: 100 μm **(C-D””, G-H””, K-L””, O-P””)** and 1 mm **(A, B)**. The nonparametric Mann–Whitney U test was used for statistical analysis, with exact p- values indicated. ns ≥ 0.05.

At E12.5 in *Tie2-Cre; R26R-Pik3ca^H1047R^; R26R-eYFP* mutant embryos, as observed at E13.5, macroscopic examination revealed blood-filled patchy lesions (**Supplemental Figure 1A**). The widespread expression of eYFP across the body in mutant mice indicates a high recombination efficiency of *Tie2-Cre* (**Supplemental Figure 1A**). In sagittal tissue sections, phospho-S6 (pS6) staining, an indicator of PI3K activity, revealed enhanced signaling in the PECAM^+^ cardinal vein and surrounding Prox1^+^ lymphatic ECs (LECs) in mutant embryos compared with controls (**Supplemental Figure 1B-B”’, C-C”’**). Analysis of cellular proliferation using the Ki67 antibody showed a higher number of positive cells in the mutant cardinal vein than in the control cardinal vein (**Supplemental Figure 1B””, C””, L**). Furthermore, most ECs in the mutant cardinal vein were eYFP^+^, indicating high recombination efficiency by Cre recombinase (**Supplemental Figure 1B’’’’’, C’’’’’**).

Next, we performed morphological analyses of the pulmonary artery, aorta, and heart. No significant differences were observed in the luminal diameters of the PECAM^+^ pulmonary artery, descending aorta, or cardiac morphology, including the PECAM^+^ endocardium, when comparing mutant and control embryos (**Supplemental Figure 1B, C, D-E”, M, N**). Both control and mutant ECs in the pulmonary artery, descending aorta, and endocardium exhibited pS6 expression, with no significant differences between them (**Supplemental Figure 1F-G’’**). Similarly, the number of Ki67^+^ ECs in the pulmonary artery, descending aorta, and endocardium did not differ between the control and mutant embryos (**Supplemental Figure 1H-I’’, O-Q**). The widespread eYFP expression in ECs of the pulmonary artery, descending aorta, and endocardium confirmed the high recombination efficiency of Cre recombinase, indicating that the lack of arterial phenotypes was not due to poor recombination (**Supplemental Figure 1J-K’’**).

### Pik3ca^H1047R^ activation timing influences vascular malformation subtypes but does not fully replicate human disease

To investigate phenotypic differences based on the timing of Pik3ca^H1047R^ expression, we crossed tamoxifen-inducible pan-endothelial *CDH5-CreERT2* mice with *R26R-Pik3ca^H1047R^* mice and analyzed the embryos at E16.5 or E17.5, focusing on the tongue, neck, liver, skin, and mesentery regions, which are frequently affected by vascular malformations in humans. When tamoxifen was administered to pregnant mice at E9.5 and analyzed at E16.5, *CDH5- CreERT2; R26R-Pik3ca^H1047R^* mutant embryos exhibited widespread, abnormal blood-filled vasculatures throughout the body (**Figure 2A, H**). In sagittal sections, the mutant mice showed an increase in slightly dilated, blood-filled PECAM^+^/Prox1^+^/VEGFR3^+^ lymphatic vessels in the tongue (**Figure 2B-C’, I-J’**). In mutant mice, the number of blood-filled PECAM^+^/Prox1^+^/VEGFR3^+^ lymphatic vessels surrounding the dilated jugular vein increased (**Figure 2D, D’, K, K’, AC**), and dilated PECAM^+^/Prox1^-^/partially VEGFR3^+^ blood vessels were observed in the liver (**Figure 2E, E’, L, L’**). Additionally, in the dorsal skin and mesentery, mutant mice exhibited an increase in dilated PECAM^+^/Prox1^-^/VEGFR3^-^ blood vessels and blood-filled PECAM^+^/Prox1^+^/VEGFR3^+^ lymphatic vessels (**Figure 2F-G’, M-N’. AC**). Following tamoxifen administration at E12.5, blood-filled vessels were less prominent, though notable edema was observed in mutant embryos (**Figure 2O, V**). In mutant embryos, PECAM^+^/Prox1^-^/partially VEGFR3^+^ enlarged blood vessels were increased in the tongue (**Figure 2P-Q’, W-X’, AD**). No significant differences were found in the PECAM^+^/Prox1^+^/VEGFR3^+^ lymphatic vessels in the neck, but mild dilation was observed in the PECAM^+^/Prox1^-^/VEGFR3^-^ jugular vein in mutant embryos compared with controls (**Figure 2R, R’, Y, Y’, AD**), and although the dilation was less extensive than at E9.5, dilated PECAM^+^/Prox1^-^/partially VEGFR3^+^ blood vessels were observed in the liver (**Figure 2S, S’, Z, Z’**). Slightly dilated, blood-filled PECAM^+^/Prox1^+^/VEGFR3^+^ lymphatic vessels were seen in the mesentery and skin of mutant mice, though the dilation was less pronounced than that observed at E9.5 (**Figure 2T-U’, AA-AB’, AD**). Mutants showed less pronounced edema when tamoxifen was administered to pregnant mice at E15.5 and analyzed at E17.5 than administration at E9.5 or E12.5, with only a few blood-filled vessels observed (**Supplemental Figure 2A**). In mutant embryos, dilated PECAM^+^/Prox1^-^/VEGFR3^+^ blood vessels and PECAM^-^/Prox1^+^/VEGFR3^+^ lymphatic vessels proliferated at the tip of the tongue (**Supplemental Figure 2B-B”, G-G”**). Blood-filled, dilated PECAM^+^/partially Prox1^+^/partially VEGFR3^+^ lymphatic vessels were observed in the skin and mesentery (**Supplemental Figure 2C-C”, H-H”, E-E”, J-J”**). Additionally, dilated PECAM^+^/Prox1^-^/VEGFR3^-^ blood vessels were observed in the liver of mutant mice (**Supplemental Figure 2F- F”, K-K’’**), but no significant differences were found in the neck region compared with controls (**Supplemental Figure 2D-D”, I-I’’**).

**Figure 2.**
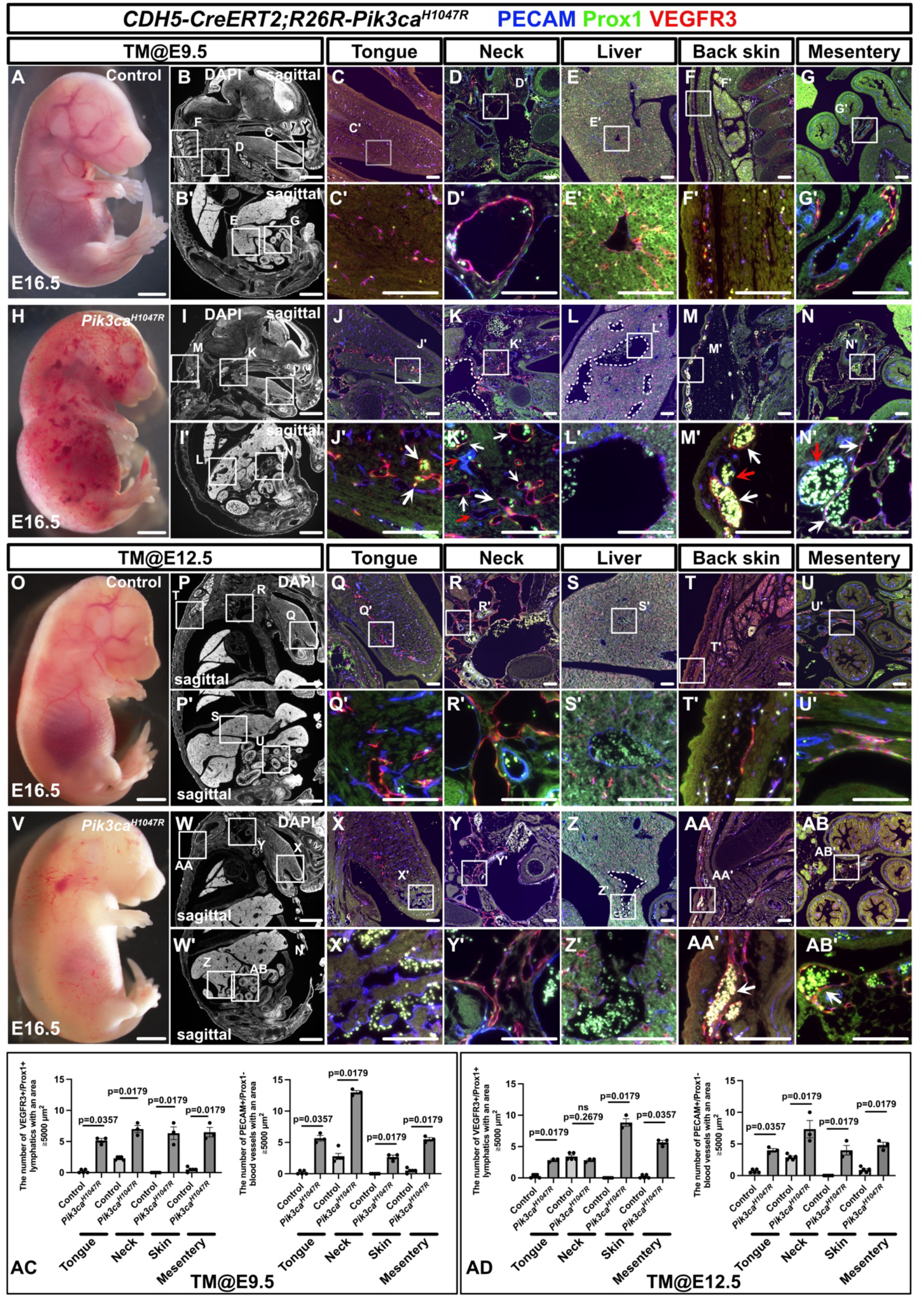
Pik3ca^H1047R^ expression timing influences the location of vascular malformations but does not fully recapitulate human pathology. **(A, H)** Gross morphology of the control and *CDH5-CreERT2; R26R-Pik3ca^H1047R^*mutant embryos at E16.5, after tamoxifen administration at E9.5. **(B-G’, I-N’)** Immunostaining of sagittal sections with the indicated antibodies. **(B, B’, I, I’)** DAPI-stained sections of the same specimen. **(D, D’, K, K’)** In the neck region of mutant embryos, dilated jugular veins are visible **(K, white dotted lines)**. Compared with controls, smaller blood-filled PECAM^+^/Prox1^+^/VEGFR3^+^ lymphatic vessels **(K’, white arrows)** and PECAM^+^/Prox1^-^ /VEGFR3^-^ blood vessels **(K’, red arrows)** are also seen. **(E, E’, L, L’)** In the liver, dilated PECAM^+^/Prox1^-^/partially VEGFR3^+^ blood vessels are observed (**L, white dotted lines**). The number of PECAM^+^/Prox1^-^/partially VEGFR3^+^ blood vessels with an area ≥ 20,000 μm^2^ in the liver is 0 ± 0 (median ± SEM) (n = 5) in controls and 3 ± 0.33 (median ± SEM) (n = 3) in mutants (p = 0.0179). **(F-G’, M-N’)** Similar observations are made in the skin and mesentery, where blood-filled PECAM^+^/Prox1^+^/VEGFR3^+^ lymphatic vessels (**M’, N’, white arrows**) and dilated PECAM^+^/Prox1^-^/VEGFR3^-^ blood vessels **(M’, N’, red arrows)** are seen. **(O, V)** Gross morphology of the control and *CDH5-CreERT2; R26R-Pik3ca^H1047R^* mutant embryos at E16.5, after tamoxifen administration at E12.5. Compared with embryos treated at E9.5, blood-filled dilated vessels are less prominent, but generalized edema is more severe. **(P, P’, W, W’)** DAPI- stained sections of the same specimen. **(P-U’, W-AB’)** Immunostaining of sagittal sections with the indicated antibodies. **(S, S’, Z, Z’)** In the liver, dilated PECAM^+^/Prox1^-^/partially VEGFR3^+^ blood vessels are observed **(Z, Z’, white dotted lines)**. The number of PECAM^+^ vessels with an area ≥ 20,000 μm^2^ in the liver is 0 ± 0 (median ± SEM) (n = 5) in controls and 3 ± 0.29 (median ± SEM) (n = 3) in mutants (p = 0.0179). **(T-U’, AA-AB’)** Dilated, blood- filled PECAM^+^/weak Prox1^+^/VEGFR3^+^ lymphatic vessels **(AA’, AB’, white arrows)** are observed in the skin and mesentery. **(AC, AD)** Statistical analysis of the tongue, neck, skin, and mesentery. Scale bars: 100 μm **(C-G’, J-N’, Q-U’, X-AB’)**, 1 mm **(B, B’, I, I’, P, P’, W, W’)**, and 2 mm **(A, H, O, V)**. The nonparametric Mann–Whitney U test was used for statistical analysis, with exact p-values indicated. ns ≥ 0.05.

These results indicate that even though the timing of Pik3ca^H1047R^ activation in ECs partially influences the location and extent of vascular malformations, it does not fully explain their localized nature in humans. In general, later induction of Pik3ca^H1047R^ leads to a more restricted anatomical distribution and fewer affected vessels than earlier induction.

### Expression of Pik3ca^H1047R^ in the CPM replicates the anatomical features of human vascular malformations

The distribution of lymphatic and blood vessels derived from the CPM aligns with the common sites of refractory vascular malformations in humans^33^. Therefore, we hypothesized that expressing Pik3ca^H1047R^ in the CPM could generate a mouse model that mimics human vascular malformations. To test this hypothesis, we crossed *Isl1-Cre;R26R-eYFP* mice, which express Cre recombinase under the control of the *Isl1* promoter and label CPM derivatives, with *R26R- Pik3ca^H1047R^* mice. Embryos of the resulting *Isl1-Cre;R26R-Pik3ca^H1047R^;R26R-eYFP* mice were analyzed at E11.5. In the mutant embryos, blood-filled vessels were observed in the first and second branchial arches, but no other significant external differences from the controls were noted (**Figure 3A**). eYFP expression was observed in the pharyngeal mesodermal region (**Figure 3A**). Sagittal sections of mutant embryos revealed enlarged PECAM^+^/Prox1^-^

**Figure 3.**
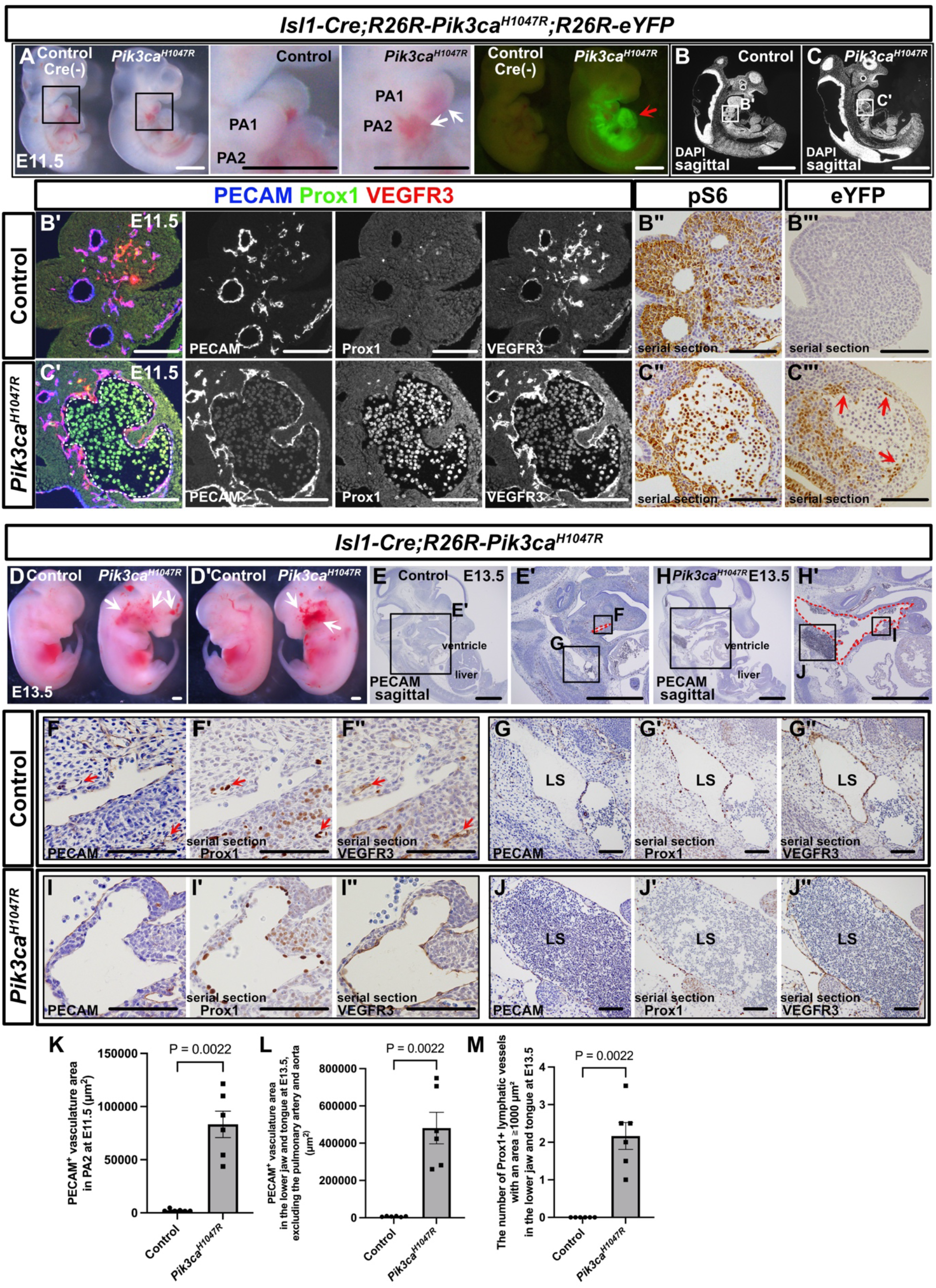
Pik3ca^H1047R^ expression in *Isl1+* CPM leads to vascular malformations confined to the head and neck region. **(A)** Gross morphology of the control and *Isl1-Cre;R26R-Pik3ca^H1047R^;R26R-eYFP* mutant embryos at E11.5. Mutant mice show widespread blood-filled, dilated vessels in the first and second pharyngeal arches, along with eYFP expression in the pharyngeal mesoderm (red arrows) **(B-C”’)** Immunostaining of sagittal sections with the indicated antibodies. Mutant mice exhibit dilated, blood-filled PECAM^+^/Prox1^-^/partially VEGFR3^+^ blood vessels in the second pharyngeal arch (**B-B’, C-C’**, white dotted lines). **(B”, B”’, C”, C”’)** Enzyme-antibody staining reveals pS6 expression in both endothelial and mesenchymal cells. eYFP, detected by GFP antibody, is expressed in ECs of dilated vessels (red arrows). **(D, D’)** Gross morphology of the control and *Isl1-Cre;R26R-Pik3ca^H1047R^*mutant embryos at E13.5. Mutant embryos display extensive blood-filled, dilated vessels on both the left and right sides of the head and neck region (white arrows). **(E-J”)** Enzyme-antibody staining of sagittal sections. In mutants, large PECAM^+^ vessels extend from the lower jaw to the jugular vein (**E-H’**, red dotted lines). Controls show small partially PECAM^+^/Prox1^+^/VEGFR3^+^ lymphatic vessels in the lower jaw, and mutants exhibit enlarged PECAM^+^/Prox1^+^/VEGFR3^+^ lymphatic vessels and blood-filled lymph sacs (**F-J”**, red arrows). **(K-M)** Statistical analysis. Each dot represents a value from one sample. Scale bars: 100 μm **(B’-C”’, F-J”)** and 1 mm **(A-C, D, D’, E, E’, H, H’)**. The nonparametric Mann–Whitney U test was used for statistical analysis.

/VEGFR3^+^ blood vessels, predominantly in the first and second branchial arches (**Figure 3B- B’, 3C-C’, K**). pS6 expression was detected in both mesenchymal cells and ECs within the branchial arches of control and mutant embryos (**Figure 3B”, C”**). Additionally, eYFP was present not only in ECs but also in mesenchymal cells within the branchial arches (**Figure 3B”’, C”’**). Similar expanded PECAM^+^/Prox1^-^/partially VEGFR3^+^ blood vessels were observed in the PA1-PA2 region of mutant embryos when *Mef2c-AHF-Cre* mice, another line that labels CPM, were crossed with *R26R-Pik3ca^H1047R^*and analyzed at E11.5 (**Supplemental Figure 3A-D”’**). In *Isl1-Cre; R26R-Pik3ca^H1047R^* embryos at E13.5, expanded vasculature was observed from the head to the neck (**Figure 3D, D’**). Sagittal sections revealed dilated vessels extending from the mandible to the tongue, along with expanded lymphatic vessels and large, blood-filled lymph sacs in mutant embryos (**Figure 3E-J”, L, M**). Next, *Isl1-CreERT2; R26R- Pik3ca^H1047R^* embryos were analyzed at E13.5 after tamoxifen administration to pregnant mice at E8.5. No significant external differences were observed compared with controls (**Supplemental Figure 3E**). eYFP expression was detected along the lower jaw, neck, and genital regions (**Supplemental Figure 3E’**). Histological sections revealed irregular, dilated PECAM^+^/Prox1^+^/VEGFR3^+^ lymphatic vessels at the junction between the tongue and lower jaw (**Supplemental Figure 3F-F”’, 3G-G”’**). eYFP^+^ cells were confirmed to contribute to ECs of these dilated lymphatic vessels (**Supplemental Figure 3F””, 3G””**).

In *Pax3-CreERT2; R26R-Pik3ca^H1047R^* embryos, which label the paraxial mesoderm (the origin of LECs in the lower body^36^), tamoxifen was administered to pregnant mice at E9.0. Upon analysis at E14.0, no vascular malformations were observed in the head or neck regions (**Supplemental Figure 3H, K**). However, mutant embryos displayed dilated, blood-filled PECAM^+^/partially Prox1^+^/VEGFR3^+^ abnormal vessels around the spine (**Supplemental Figure 3I-J”, L-M”)**. In *Myf5-CreERT2; R26R-tdTomato*, which labels satellite cells in the musculature, crossed with *R26R-Pik3ca^H1047R^*, tamoxifen was administered to pregnant mice at E9.5. Upon analysis at E14.5, no external differences were observed between the control and mutant embryos (**Supplemental Figure 3N**). tdTomato expression was observed in the face and limbs (**Supplemental Figure 3N’**). Histological analysis showed no major changes in sagittal sections, and tdTomato^+^ cells contributed to tongue skeletal muscle without hypertrophy or hyperplasia in mutants. Additionally, PECAM^+^ vasculature showed no significant differences between the controls and mutants (**Supplemental Figure 3O-P’’**).

These findings suggest that ECs are susceptible to the effects of the *Pik3ca^H1047R^*mutation and that their cellular origin dictates the anatomical site where vascular malformations develop.

### Single-cell RNA sequencing analysis elucidates the endothelial cell differentiation process from the CPM

To clarify the causes of diverse vascular malformations in the head and neck and establish a foundation for understanding how ECs differentiate from CPM, we re-analyzed previously published scRNA-seq data^37^. Specifically, we analyzed the differentiation process of *Mesp1^+^* CPM cells into ECs. UMAPs were generated from *Mesp1^+^* CPM populations at E8.0, E8.25, E9.5, and E10.5 (**Supplemental Figure 4A, B**), identifying EC clusters expressing various markers, such as *Pecam1, Kdr, Cdh5, Tek*, and *Etv2* (Clusters 7 and 15), whereas *Isl1* expression was not highly detected (**Supplemental Figure 4A-D, Supplemental Data 1**). Further subclustering of these two clusters revealed eight subclusters, labeled 0 through 7 (**Supplemental Figure 4E-H, Supplemental Data 1**). Additionally, RNA velocity and trajectory inference analyses were performed to predict the differentiation pathways of ECs (**Supplemental Figure 4I**). The results suggested a lineage progression from *Isl1^+^*undifferentiated mesodermal cells to *Npr3*^+^ endocardial cells^38^ or *Etv2*^+^ EC progenitor cells (PCs) (**Supplemental Figure 4I, J**). *Prox1*^+^ LEC PCs appear to arise directly from *Etv2*^+^ EC progenitors, bypassing the venous stage, whereas *Aplnr^+^* venous ECs and *Efnb2*^+^ arterial ECs likely develop from shared arterial/venous PCs (A/V PCs) (**Supplemental Figure 4J**).

To further investigate the gene expression changes that drive the differentiation of immature ECs from undifferentiated mesodermal cells, we conducted a more detailed analysis using only the data from E8.0 and E8.25. When projected onto UMAP, the data revealed 15 distinct clusters (**Supplemental Figure 5A-C, Supplemental Data 2**). Widespread expression of the CPM markers *Isl1* and *Wnt5a* was observed (**Supplemental Figure 5D**). We focused on clusters 4, 11, 12, and 3, which contained immature ECs, as indicated by markers such as *Etv2, Kdr, Flt4*, and *Pecam1* (**Supplemental Figure 5C, D, Supplemental Data 2**). Subclustering analysis of these clusters revealed seven subclusters, labeled 0 through 6 (**Supplemental Figure 5E-H, Supplemental Data 2**). RNA velocity and trajectory analyses revealed early expression of *Isl1* and *Wnt5a*, which decreased as EC differentiation progressed. This was followed by the expression of key endothelial differentiation markers, *Etv2* and *Kdr*, with subsequent expression of more mature markers such as *Flt4* and *Pecam1* (**Supplemental Figure 5E-J**).

### *Pik3ca*-driven vascular malformations are associated with hypoxia-mediated metabolic changes and Vegf-a expression

To investigate how Pik3ca^H1047R^ expression in the CPM affects ECs and drives gene expression changes contributing to vascular malformations, we conducted scRNA-seq analysis on *Isl1- Cre; R26R-Pik3ca^H1047R^; R26R-eYFP* mutant embryos and *Isl1-Cre; R26R-eYFP* controls at E13.5, the stage when lymphatic vessels form lumens in the head and neck. After FACS sorting of eYFP^+^ cells, the sorted cells were analyzed (**Figure 4A, Supplemental Figure 6A**). UMAP analysis identified 15 clusters, with cluster 1 defined as ECs (**Figure 4B, C, Supplemental Data 3**). In mutant mice, the proportion of ECs increased compared with controls, while the proportion of mesenchymal cells decreased (**Figure 4B, D, Supplemental Data 3**). Enrichment analysis highlighted significant changes in hypoxia and glycolysis pathways, which were observed not only in ECs but also in cardiomyocytes, pharyngeal arch muscles, neurons, epithelium, and mesenchyme (**Figure 4E, Supplemental Figure 6B, Supplemental Data 3, 4**). Bulk RNA-seq analysis of HUVECs expressing PIK3CA^H1047R^ and ECs from patients with *PIK3CA* gain-of-function mutations also showed elevated hypoxia and glycolysis pathways (**Figure 4E, Supplemental Figure 6C, Supplemental Data 3, 4**). The distribution of cells in the G1, G2/M, and S phases was similar across cell types, except for neurons, where over 80% were in the G1 phase (**Supplemental Figure 6D, Supplemental Data 4**). Volcano plots for cluster 1 ECs highlighted genes associated with hypoxia and glycolysis (**Figure 4F, G, Supplemental Data 3**). Several genes involved in hypoxia and glycolysis, including *Vegf- a, Flt1, Adm, Slc2a1, Pgk1, Tip1*, and *Pkm*, were upregulated in mutant embryo ECs. Correspondingly, hypoxia-inducible factor (*Hif*) stabilizing genes, such as *Nfe2l2, Atf4, Ddit,* and *Pdk1*, were also upregulated in mutant EC clusters. Additionally, Hsp family members (*Hsp90b1, Hsp90aa1, Hspa1a, Hspa1b, Hspa5*), possibly associated with Hif stabilization, also showed increased expression (**Figure 4F, Supplemental Data 3**). However, the expression levels of *Vegfb, Vegfc, Vegfd, Kdr*, and *Flt4* were not significantly different between the control and mutant mice (**Figure 4G, Supplemental Figure 6E, Supplemental Data 3, 4**).

**Figure 4.**
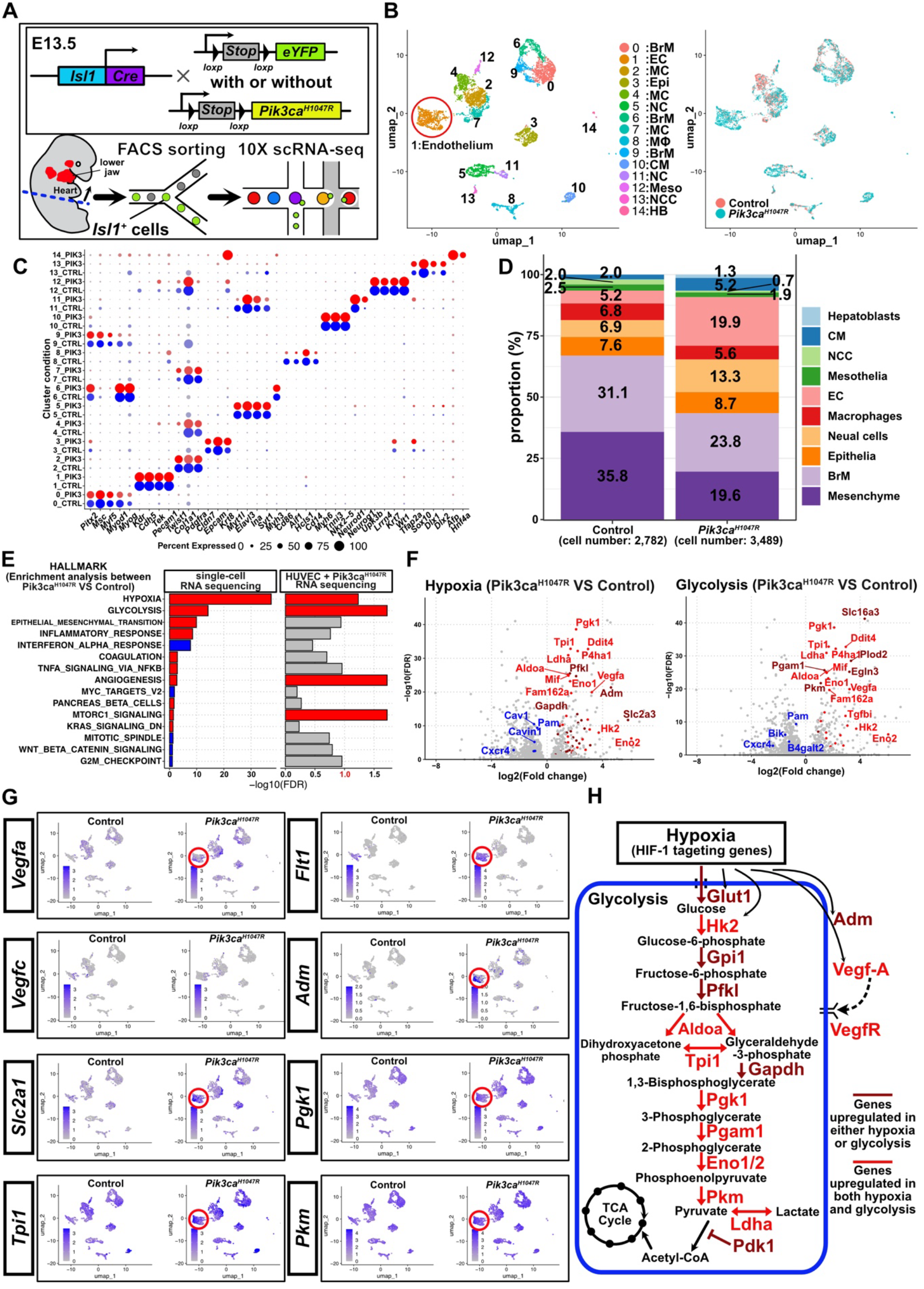
Pik3ca^H1047R^ induces hypoxia-mediated metabolic changes in endothelial cells. **(A)** Schematic overview of the cell isolation conducted for scRNA-seq. Metrics for E13.5 *Isl1- Cre; R26R-eYFP* and *Isl1-Cre; R26R-eYFP; R26R-Pik3ca^H1047R^* samples showed 65.2% reads in cells for *Isl1-Cre; R26R-eYFP* (mean 52,351 reads/cell, median 3,850 genes, 4,505 cells estimated) and 67.9% reads for *Isl1-Cre; R26R-eYFP; R26R-Pik3ca^H1047R^* (mean 43,227 reads/cell, median 3,529 genes, 5,357 cells). Filtering based on gene count (2,000–9,000) and mitochondrial content (< 10%) resulted in 2,782 and 3,489 cells for *Isl1-Cre; R26R-eYFP* and *Isl1-Cre; R26R-eYFP; R26R-Pik3ca^H1047R^*, respectively, with 6,271 cells included in downstream analyses. **(B)** Left: UMAP plot showing color-coded clusters (0–14). 0: branchial muscles (BrM); 1: endothelial cells (EC); 2: mesenchymal cells (MC); 3: epithelia (Epi); 4: MC; 5: neural cells (NC); 6: BrM; 7: MC; 8: macrophages (Mφ); 9: BrM; 10: cardiomyocytes (CM); 11: NC; 12: mesothelia (Meso); 13: neural crest cells (NCC); and 14: hepatoblasts (HB). Right: UMAP plot color-coded by condition. Pink represents the control, and light blue represents mutant cells. **(C)** Heatmap showing the average gene expression of marker genes for each cluster by condition. **(D)** Cell type proportions. **(E)** Comparison of enrichment analysis between EC clusters from scRNA-seq and bulk RNA-seq of HUVECs. The left bar graph shows the top 15 significantly altered Hallmark gene sets in EC clusters from scRNA-seq using ssGSEA (escape R package). The right bar graph shows the re-analysis of public bulk RNA-seq from PIK3CA^H1047R^-transfected HUVECs. Red bars represent significantly upregulated Hallmark gene sets in mutants (FDR < 0.1), and gray bars indicate non-significant sets. **(F)** Left: Volcano plot showing hypoxia-related genes (red: mutant upregulated, blue: control upregulated, dark red: hypoxia only, bright red: hypoxia and glycolysis). Right: Glycolysis-related genes with the same color scheme. Differential expression is defined as FDR < 0.05 and fold change >1.5. **(G)** UMAP plot showing expression levels of genes related to hypoxia or glycolysis. **(H)** Schematic showing upregulated genes in scRNA-seq. Bright red indicates genes involved in both hypoxia and glycolysis, while dark red represents genes specifically upregulated in either the hypoxia or glycolysis pathway.

In summary, mutant ECs showed increased expression of genes involved in hypoxia and glycolysis, including Hif target genes, such as *Vegf-a* and *Adm*, which stimulate angiogenesis and lymphangiogenesis. Glycolytic enzymes were also upregulated. Notably, *Pdk1*, which inhibits pyruvate conversion to acetyl-CoA, and lactate dehydrogenase A (*Ldha*) were elevated, indicating a shift toward anaerobic metabolism, similar to that seen in malignant tumors and actively proliferating cells (**Figure 4H, Supplemental Data 3**).

### Elevated HIF-1α and VEGF-A expression is observed in *PIK3CA*-mutated human vascular malformations

To determine whether the signals identified by scRNA-seq were altered in human vascular malformation samples, we focused on HIF-1α and VEGF-A and confirmed protein expression by immunostaining. In control human epicardial tissue, PECAM^-^/podoplanin^+^ LECs and PECAM^+^/podoplanin^-^ venous ECs did not express VEGF-A or HIF-1α. However, in PECAM^+^/podoplanin^-^ arterial ECs, mild expression of VEGF-A was observed in both the vessel wall and ECs, and HIF-1α was detected in these locations (**Figure 5A-C””’**). HIF-1α and VEGF-A were not detected in negative controls (**Figure 5A””’, B””’, C””’**). In LM samples with the *PIK3CA^H1047R^* mutation, particularly in clinically microcystic cases with extensive fibrosis and chronic inflammatory cell infiltration, irregularly structured PECAM^+^/partially podoplanin^+^ abnormal lymphatic vessels within fibrotic tissue showed partial expression of VEGF-A and HIF-1α (**Figure 5D-I**). Nuclear expression of HIF-1α was observed (**Figure 5I**). pS6 expression was detected in a subset of malformed ECs (**Figure 5J**). No signal was detected in the negative control (**Figure 5K**). A similar expression pattern was observed in macrocystic lymphatic vessels (**Supplemental Figure 7A-H**). Comparable findings of HIF-1α, VEGF-A, and pS6 expression were noted in venous malformation patients with the same mutation (**Figure 5L-S**) and in LM patients with *PIK3CA^E542K^* and *PIK3CA^E545K^* mutations (**Supplemental Figure 7I-X**).

**Figure 5.**
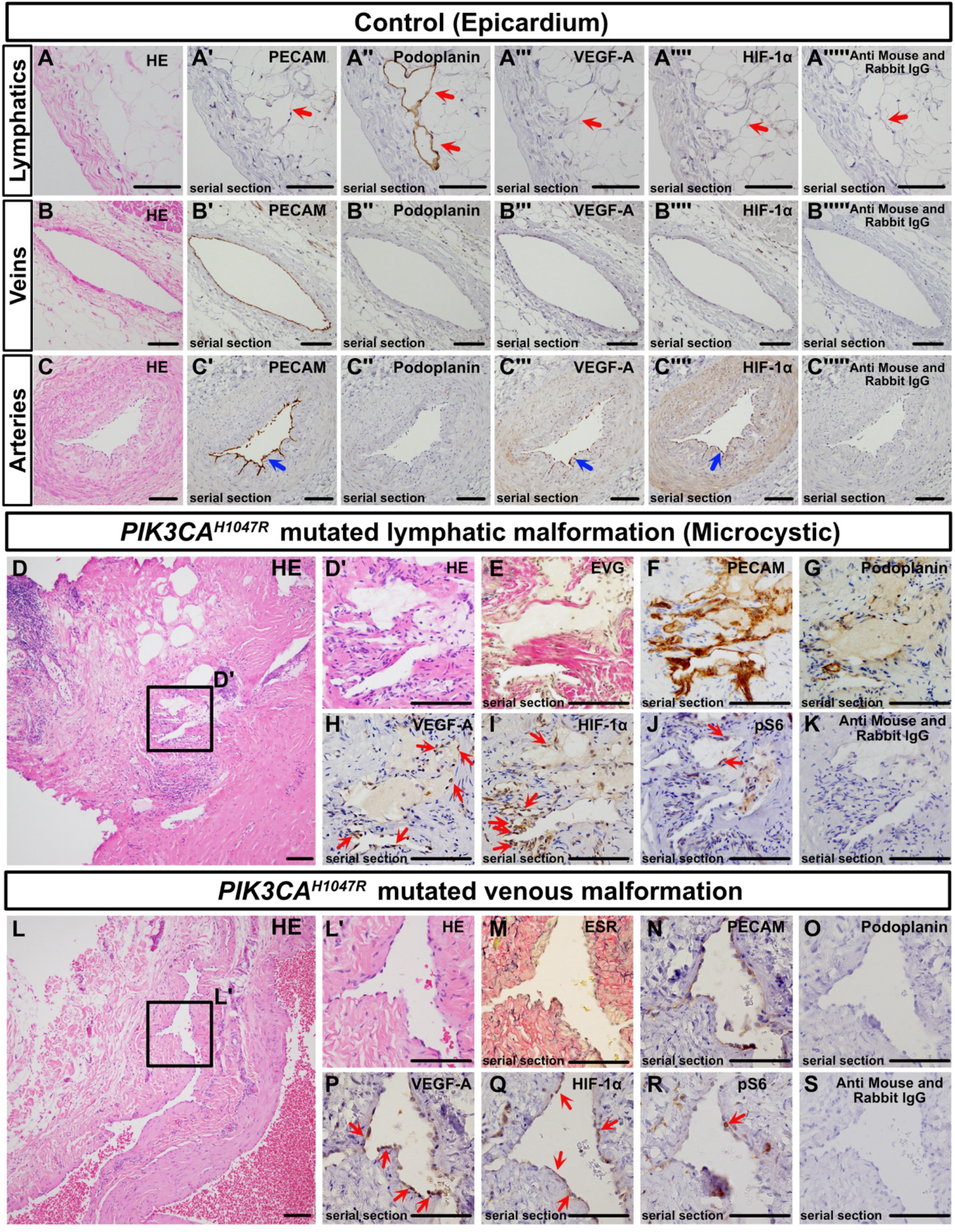
HIF-1α and VEGF-A expression in endothelial cells of malformed lymphatic vessels and veins in human patient samples. **(A-S)** Hematoxylin and eosin (HE) staining, special staining [elastica van Gieson (EVG), elastica Sirius red (ESR)—where collagen fibers appear red, elastic fibers black, and muscle tissue yellow], and immunohistochemistry using the indicated antibodies. No signal was detected in negative controls using only secondary antibodies **(A””’, B””’, C””’, K, S)**. **(A- A””’)** Lymphatic vessels (red arrows) on the epicardium, characterized as PECAM^-^/podoplanin^+^, do not express HIF-1α or VEGF-A. **(B-B””’)** Similarly, PECAM^+^/podoplanin^-^ veins show no expression of these markers. **(C-C””’)** In PECAM^+^/podoplanon^-^ muscular arteries, both VEGF-A and HIF-1α are expressed in the arterial walls and endothelium (blue arrows). **(D-K)** Microcystic-type lymphatic malformations with the *PIK3CA^H1047R^* mutation. Among the fibrotic tissue background **(D)**, small, irregular PECAM-overexpressing vessels **(E)**, partially podoplanin^+^ **(G)** lymphatic vessels are found. These vessels express VEGF-A and HIF-1α, and partially express pS6 (**H-J,** red arrows). **(L-S)** Venous malformation samples with the *PIK3CA^H1047R^* mutation. These vessels have blood-filled lumens and thicker walls than those in lymphatic malformations. Similar to the lymphatic vessels, the malformed venous endothelium expresses VEGF-A and HIF-1α (**P-Q**, red arrows). Scale bars: 100 μm **(A-C””’, D’-K, L’-S)** and 1 mm **(D, L)**.

## Hif-1α and Vegf-A inhibitors suppress the progression of vascular malformations

We next examined whether the administration of Hif-1α and Vegf-A inhibitors could potentially treat vascular malformations. Tamoxifen was administered to 3–4-week-old *CDH5- CreERT2;R26R-Pik3ca^H1047R^*mice to induce mutations in the dorsal skin. Bevacizumab, a Vegf-a inhibitor, LW6, a Hif-1α inhibitor, and rapamycin, an mTOR inhibitor, were topically applied, and the effects were analyzed (**Figure 6A**). Both bevacizumab and LW6 reduced the visible swelling in the dorsal skin, whereas the difference between drug-treated and control groups was less pronounced with rapamycin (**Figure 6B**). Compared with normal skin, tamoxifen treatment resulted in enlarged, disorganized PECAM^+^ vasculatures extending from the dermis to the subcutaneous tissue, with additional signs of inflammatory cell infiltration and fibrosis in the background (**Figure 6C, D**). The number of PECAM^+^ vasculatures significantly decreased in the bevacizumab, LW6, and rapamycin groups, with bevacizumab and LW6 showing superior efficacy (**Figure 6D**). Larger PECAM^+^ vasculatures were also reduced with bevacizumab and LW6 treatment **(Figure 6D**). A similar pattern was observed for VEGFR3^+^ lymphatics, though rapamycin showed limited effectiveness. Bevacizumab and LW6 were particularly effective in reducing malformed vasculatures, with LW6 demonstrating the most pronounced effect (**Figure 6C, E**).

**Figure 6.**
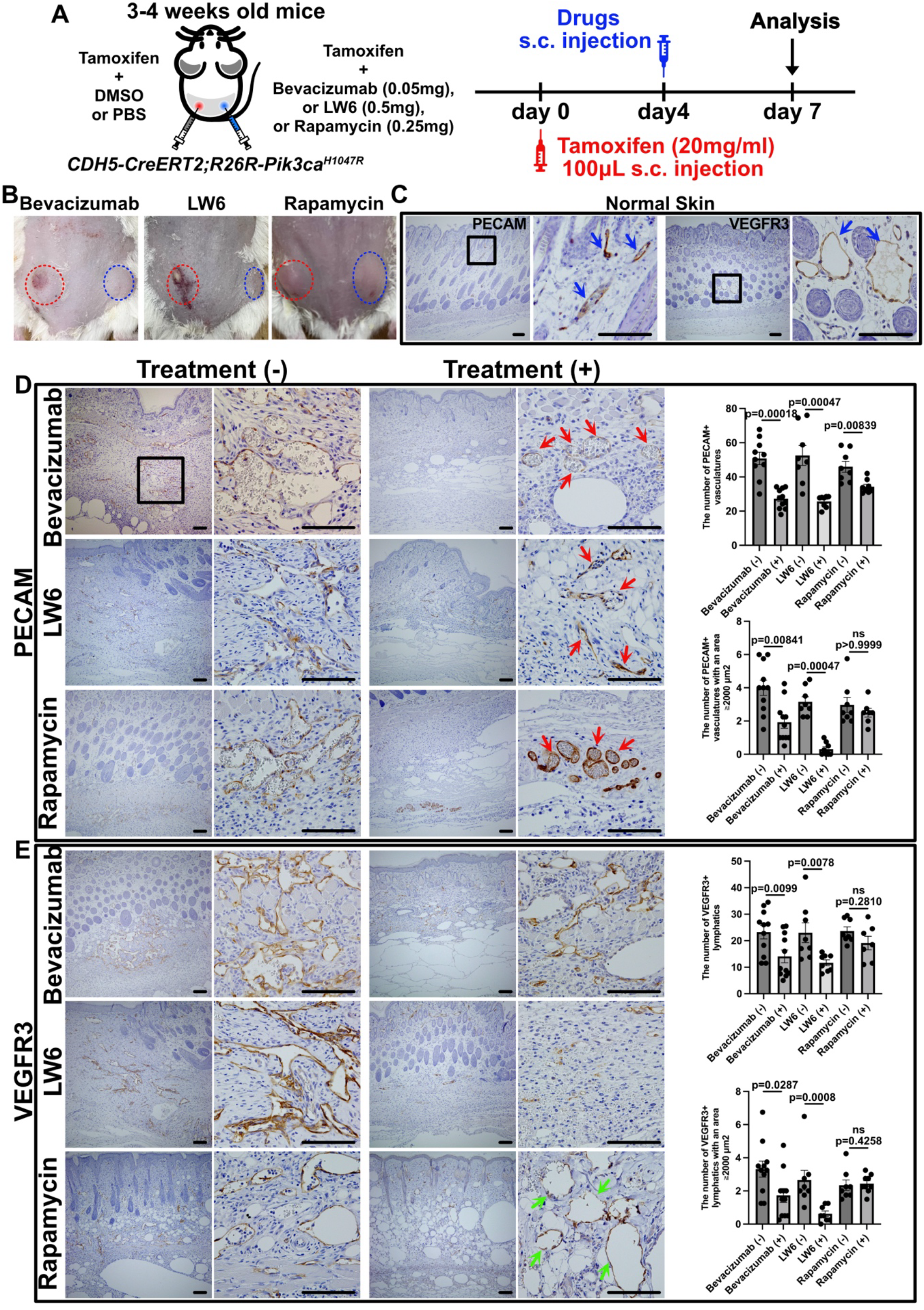
HIF-1α and VEGF-A inhibitors significantly suppress vascular malformations in the dorsal skin of mice. (**A**) Experimental design for inducing and treating progressive vascular malformations with bevacizumab, LW6, rapamycin, or control (DMSO for LW6 and rapamycin; PBS for bevacizumab). (**B**) Macroscopic effects of treatment: the left panel shows the control group (red dotted line), and the right panel shows the treatment group (blue dotted line). (**C**) Normal dorsal skin, displaying PECAM^+^ blood vessels and VEGFR3^+^ lymphatic vessels (blue arrows). (**D, E**) Sections were prepared vertically from the epidermis to the dermis. Immunohistochemistry was performed using the indicated antibodies. The treatment groups exhibited a reduction in PECAM^+^ vessels (red arrows). VEGFR3^+^ lymphatic vessels were markedly reduced in the LW6 group, with a similar reduction observed in the bevacizumab group. However, the rapamycin-treated group retained noticeably enlarged blood vessels (green arrows). Each dot represents data from an individual sample. The white, vacuole-like structures in the image are oil droplets from the tamoxifen injection. Scale bars: 100 μm **(C, D, E)**. Statistical analysis was performed using the nonparametric Mann–Whitney U test. ns ≥ 0.05.

## Discussion

The common *PIK3CA*-activating mutation H1047R, which is linked to both VMs and LMs, as well as its association with cancer and PROS, has been well established^9,11,13,15,16,39^. However, the anatomical factors contributing to the complexity of treating vascular malformations and the changes in downstream signaling pathways remain insufficiently understood. Our analysis showed that Pik3ca^H1047R^ expression in the CPM recapitulates the head and neck refractory vascular malformations. From scRNA-seq, we identified the significant upregulation of genes related to hypoxia and glycolysis in Pik3ca^H1047R^-expressing ECs. Consistent with these findings, elevated HIF-1α and VEGF-A expression was observed in human samples of both LMs and VMs with various *PIK3CA* mutations. Additionally, topical application of Hif-1α and Vegf-a inhibitors in a *CDH5-CreERT2; R26R-Pik3ca^H1047R^*skin model led to a significant reduction in lesion formation. In summary, our findings reveal hypoxia-driven molecular signaling changes in ECs harboring this mutation.

Our findings suggest that embryological cellular origins may play a crucial role in anatomically characterizing refractory vascular malformations. Although the CPM contributes to connective tissues, skeletal muscles, and cardiomyocytes, our study only reproduced vascular malformations, in contrast to the overgrowth observed in conditions like PROS. Similarly, Castillo et al. found that early expression of Pik3ca^H1047R^ in mesodermal cells resulted in VMs without organomegaly^16^. In contrast, human PROS often presents with organomegaly during embryonic development^40^. These discrepancies may stem from differences in the types of *PIK3CA* mutations^41^ or tissue-specific sensitivities. For instance, *Pik3ca^H1047R^* mutations replicate brain organomegaly in mouse models of PROS^42^. Moreover, *PIK3CA* mutations have not been associated with arterial malformations^43,44^. Given that PI3K/AKT signaling is critical for arterial specification^45,46^, arteries, which already have sufficient PI3K/AKT signaling, may be less affected by these mutations^47^. Consistent with this, no significant differences in pS6 expression were observed between control and mutant aortic, pulmonary arterial, and endocardial ECs in *Tie2-Cre;R26R-Pik3ca^H1047R^*embryos (**Supplemental Figure 2**).

By analyzing the differentiation of *Mesp1^+^* CPM cells^37^, we explored how ECs in the head and neck region change their gene expression and at what stage their fate is determined. Our analysis showed that *Isl1* expression decreases at the mRNA level during early embryogenesis (E8.0–E8.25), followed by the expression of *Etv2, Kdr*, and eventually *Pecam*, marking the maturation into ECs. Additionally, our findings suggest that lymphatic vessels may arise directly from a common progenitor shared with veins, rather than through venous endothelium (**Supplemental Figures 4, 5**). Given that *Isl1* expression disappears at a very early stage and contributes to endothelial differentiation, experiments using *Isl1-Cre* or *Isl1-CreERT2* mice cannot clearly distinguish between LMs, VMs, and capillary malformations. Therefore, the diverse types of vascular malformations in the head and neck, including mixed venous-lymphatic and capillary malformations, as well as the macro and microcystic subtypes of LMs, cannot be fully explained by this study alone. Additionally, the timeline of development significantly differs between humans and mice. In humans, lymphatic vessels in the trunk are formed within approximately one week, whereas it takes approximately four weeks for lymphatic vessels in the head and neck to establish a lumen structure^34^. A deeper understanding of Pik3ca^H1047R^’s impact on cell differentiation requires further investigation.

Our scRNA-seq analysis revealed a notable increase in EC numbers and significant changes in hypoxia- and glycolysis-associated gene expression. Similar shifts were observed in other cell types, but they did not result in phenotypic changes, suggesting that ECs are particularly susceptible to these molecular alterations. We focused on Vegf-a, a key regulator of EC proliferation and downstream target of Hif-1α, which likely drives both cell-autonomous and non-cell-autonomous effects on ECs. The increased expression of glycolytic enzymes, including *Ldha*, further suggests a shift toward anaerobic metabolism (Warburg effect). Previous studies have also shown that the stabilization of HIF-1α in cancerous tumors and pulmonary arterial smooth muscle cells is regulated by the PI3K/AKT signaling pathway^48,49^. Additionally, the Warburg effect, driven by HIF-1α, promotes EC proliferation and migration^50,51^. Our findings indicate that vascular malformations may share mechanistic similarities with cancer. Notably, mTOR inhibitors^52^, commonly used in cancer therapies, have shown efficacy in reducing symptoms of vascular malformations. Likewise, HIF and VEGF inhibitors, employed in the treatment of malignant tumors, induce a significant reduction in malformed vasculature.

Our study reveals previously unrecognized mechanisms behind *PIK3CA*-driven vascular malformations and highlights the role of hypoxia-mediated signaling pathways in disease pathogenesis. Furthermore, we propose novel therapeutic strategies targeting these mechanisms. This research supports the development of new treatment approaches for intractable vascular malformations that afflict patients from childhood into adulthood.

## Materials and methods Human samples

Formalin-fixed paraffin-embedded (FFPE) tissues of residual specimens used for diagnostic purposes were employed. These surgical specimens were classified according to the classification system proposed by the International Society for the Study of Vascular Anomalies^53^. This study was approved by the Ethical Review Board of the Graduate School of Medicine, Mie University (approved number: H2024-078) and the Graduate School of Medicine, Osaka University (approved number: 17214 and K24132). Informed consent was supplemented by an opt-out provision, ensuring the participants’ autonomy to withdraw from the study at any point. The human study was conducted in compliance with Japanese regulations, following the Ethical Guidelines for Medical and Health Research Involving Human Subjects, in addition to the Declaration of Helsinki. Mutation analysis was conducted by Y.H. and H.K. following the protocols^14^. Next-generation sequencing (NGS) was performed using a custom panel, as previously described^54^. Genomic DNA was extracted from FFPE tissues using the QIAamp DNA FFPE Tissue Kit (Qiagen, Valencia, CA, USA). The gene panel was designed using SureDesign (https://earray.chem.agilent.com/suredesign, accessed Dec 10, 2023) to cover all the exons of the *PIK3CA* genes. Sequence libraries were prepared using the custom SureSelect Low-Input Target Enrichment System (Agilent Technologies Inc., Santa Clara, CA, USA) and sequenced using an Illumina MiSeq instrument (Illumina, San Diego, CA, USA). Variant calling was performed using SureCall version 4.0 (https://www.agilent.com/en/download-software-surecall, accessed Dec 10, 2023). Intron DNA, non-coding DNA, and variants with an allele frequency of less than 1% were excluded. Variants obtained by panel sequencing were confirmed by Sanger sequencing^14, 54^.

## Immunohistochemistry and special staining

For histological analyses, samples were collected and fixed in 2% paraformaldehyde overnight at 4°C, followed by storage in 70% ethanol at 4°C. HE staining and IHC were performed using 3 μm-thick paraffin-embedded sections. Sections for IHC were deparaffinized and rehydrated through a series of xylene and ethanol. For the enzyme-antibody method, endogenous peroxidase activity was blocked using 0.3% hydrogen peroxide (H2O2) in methanol for 20 minutes. For fluorescent antibody staining, to suppress autofluorescence, samples were incubated in 0.1% sodium borohydride in 0.1 M phosphate-buffered saline (PBS) for 30 minutes, then rinsed with water, and subsequently incubated for 5 minutes in 0.2 M glycine in 0.1 M PBS. Antigen retrieval was carried out using a pressure chamber with Tris-EDTA buffer (7.4 mM Tris, 1 mM EDTA-2Na, pH 9.0). The sections were immunostained using primary antibodies against PECAM (DIA-310, Dianova, 1:100, RRID:AB_2631039; M0823, DAKO, 1:100, RRID:AB_2114471), Prox1 (11-002, AngioBio, 1:100, RRID:AB_10013720; AF2727, R&D Systems, 1:100, RRID:AB_2170716), VEGFR3 (AF743, R&D Systems, 1:100, RRID:AB_355563), GFP (ab290, Abcam, 1:250, RRID:AB_303395), phospho-S6 (#2215, CST, 1:100, RRID:AB_331682), Ki67 (ab15580, Abcam, 1:250, RRID:AB_443209), D2-40 (anti-PDPN) (413151, Nichirei Biosciences, ready to use), HIF-1α (ab114977, Abcam, 1:100, RRID:AB_10900336), and VEGF-A (ab52917, Abcam, 1:100, RRID:AB_883427; ab51745, Abcam, 1:100, RRID:AB_2256948). For the enzyme-antibody method, the secondary antibody from the Histofine Simple Stain System (Nichirei Biosciences) was incubated with the slides for 1 hour. Peroxidase activity was visualized using DAB-H_2_O_2_. For fluorescent immunostaining, Alexa Fluor-conjugated secondary antibodies (Abcam, 1:400) were subsequently applied. Imaging was carried out using a Keyence BZ-X700 microscope. All images were processed using ImageJ software.

For Elastica van Gieson (EVG) staining, tissue sections were deparaffinized in xylene, rehydrated through graded ethanol, and rinsed in Milli-Q water. Sections were then treated with 1% hydrochloric acid in 70% ethanol, stained with Weigert’s resorcin-fuchsin for 40–50 minutes to stain elastic fibers dark purple-black, and counterstained with Weigert’s iron hematoxylin for 3–5 minutes. After additional rinsing, sections were stained in Van Gieson’s solution for 15 minutes, which stains collagen fibers red, muscle tissue yellow, and elastic fibers black. For Elastica Picrosirius Red (ESR) staining, tissue sections were deparaffinized, rehydrated through graded ethanol, and rinsed in Milli-Q water. Sections were treated with 1% hydrochloric acid in 70% ethanol, stained with Weigert’s resorcin-fuchsin for 40–50 minutes to visualize elastic fibers (stained dark purple-black), and counterstained with Weigert’s iron hematoxylin for 3–5 minutes. After additional rinsing, sections were stained in Picrosirius Red solution for 15 minutes, which stains collagen fibers red. In this protocol, muscle tissue is also stained yellow, and elastic fibers appear black. Reagents for special staining were purchased from Muto Pure Chemicals Co., Tokyo.

## Quantification of section immunostaining

For quantifying immunostained sections, an average of at least two 3 μm-thick sections from 20× power fields (0.55 mm^2^/field) for each anatomical region were analyzed. For sagittal sections, one HE-stained section and nine unstained sections were prepared per embryo, typically with a total of 100 sections made. The sections were centered around the position containing the ascending aorta and heart. In the case of transverse sections, the samples included all sections from the head down to where the heart was no longer visible.

## Mouse lines and treatments

The following mouse strains were used: *Tie2-Cre*^55^*, CDH5-CreERT2*^56^, *Isl1-Cre*^57^, *Mef2c- AHF-Cre*^58^, *Isl1-CreERT2*(**RRID:IMSR_JAX:029566**)^59^, *Myf5-CreERT2* (**RRID:IMSR_JAX:023342**)^60^, *Pax3-CreERT* (**RRID:IMSR_JAX:025663**)^61^, *R26R-eYFP*^62^, *R26R-tdTomato*(**RRID:IMSR_JAX:007914**)^63^, and *R26R-Pik3ca^H1047R^* (**RRID:IMSR_JAX: 016977**)^64^. Mice with the indicated RRID were purchased from The Jackson Laboratory, USA. All mice were maintained on a mixed genetic background (C57BL/6J x Crl:CD1(ICR)), both female and male mice were used for analyses, and no differences in phenotype were observed between them. The genotypes of the mice were determined by polymerase chain reaction using tail-tip or amnion DNA and the primers listed in **Supplemental Table 1**. The mice were housed in an environmentally controlled room at 23 ± 2°C, with a relative humidity level of 50%–60%, under a 12-h light:12-h dark cycle.

Embryonic stages were determined by timed mating, with the day of the appearance of a vaginal plug designated as embryonic day (E)0.5. All animal experiments were approved by the Mie University animal care and use committee and performed in accordance with institutional guidelines.

For embryo experiments, tamoxifen (20 mg/mL; Sigma Aldrich, T5648) was dissolved in corn oil, and pregnant mice were intraperitoneally administered (125 mg/kg body weight) at the indicated time points. In experiments using *CDH5-CreERT2*, tamoxifen was diluted to one-quarter strength (31.25 mg/kg body weight) in corn oil and intraperitoneally injected into pregnant mice.

## Mouse model for skin vascular malformations

For postnatal induction, 3 to 4-week-old *CDH5-CreERT2; PIk3ca^H1047R^* mice and controls were used. The dorsal region was shaved using clippers and depilatory cream (Reckitt Benckiser). Tamoxifen (20 mg/mL) was intradermally administered (100 μL) into the skin above both thighs, ensuring a localized circular swelling to confine the injection area. Four days later, the right side received one of the following treatments applied directly to the tamoxifen-treated area to target malformed vasculature: LW6 (Hif-1α inhibitor, S8441, Selleck), dissolved in DMSO (5 mg/mL), with 100 μL administered; bevacizumab (VEGF-A inhibitor, A2006, Selleck), prepared in PBS (1 mg/mL), with a single 50-μL injection; and rapamycin (mTOR inhibitor, S1039, Selleck), dissolved in DMSO (5 mg/mL), with 50 μL administered. The left side was treated with the corresponding solvent (PBS or DMSO) as a control. The dosage of the drugs was determined based on previously published literature^65–67^ and information provided on the Selleck Chemicals website. Three days after treatment, the skin was excised down to the muscle using scissors, processed for paraffin embedding, and subjected to immunohistochemistry analysis. Sections were stained for PECAM and VEGFR3. Additionally, untreated skin from the upper dorsal region was harvested and used as a normal control.

## Embryo preparation for single-cell RNA sequencing and FACS sorting

E13.5 embryos were collected by cesarean section from *Isl1-Cre;R26R-eYFP* or *Isl1- Cre;R26R-eYFP;Pik3ca^H1047R^* mice. The embryos were washed twice with 0.1 M PBS. Using a stereomicroscope, three eYFP^+^ embryos were selected per group. The upper body regions, including tissues above the liver, were dissected and finely minced using scissors in 10 ml of DMEM (Fujifilm, 041-30081) containing 10% FBS (Gibco, 10270106), 1% antibiotic-antimycotic (AB/AM) solution (Gibco, 15240062), 0.5 mg/ml DNase I (Roche, EN0521), and 1 mg/ml type II collagenase (Worthington, CLS-2). The tissue suspension was incubated at 37°C in a water bath for 30 minutes, then filtered through a 70-µm cell strainer (Falcon, 352350), followed by two passages through a 40-µm cell strainer (Corning, 431750). After staining, cells were washed with FACS buffer (PBS, 0.5% FBS, 2 mM EDTA), and the cell pellet was resuspended in 1 ml of 2 mM EDTA/PBS/0.1% BSA for FACS sorting. Cell sorting was performed using FACS Aria III (BD Biosciences). Data were analyzed using BD FACSDiva software.

## Analysis of the single-cell RNA-seq results

Libraries were prepared using the Next GEM Single Cell 5′ Library and Gel Bead Kit v2 (10x Genomics), following the manufacturer’s protocol. Raw reads were processed using 10x Genomics Cell Ranger software (v7.1.0) for demultiplexing, genome alignment (GRCm38/mm10), and feature-barcode matrix generation. Data analysis was performed using R (version 4.3.3) and the Seurat package (version 5.0.3). Quality control was conducted based on unique molecular identifiers, gene counts, and mitochondrial gene expression. Cell cycle gene expression was regressed during the integration step to avoid clustering bias. Data from multiple samples were integrated using both Anchor-based CCA (Seurat version 5.0.3) and Harmony integration (harmony package version 1.2.0), and no significant differences were found between these methods. Therefore, Anchor-based CCA was used for downstream analyses. Principal component analysis (PCA) was performed for dimensionality reduction, followed by UMAP for cluster visualization. To address cell cycle heterogeneity, cell cycle phase scores were calculated using known marker genes^68^, and the cell cycle effects were regressed using the *CellCycleScoring* and *ScaleData* functions.

To annotate each cluster, conserved marker genes between conditions were detected using the *FindConservedMarkers* function from the Seurat package (FDR < 0.05, log2-fold change >1).

For differentially expressed gene analysis, the *FindMarkers* function from the Seurat package was used (FDR < 0.05, fold change > 1.5). For enrichment analysis, we utilized Hallmark gene sets from the MSigDB Collections, applying two methods: over-representation analysis (Enrichr web tool^69^) and single-sample Gene Set Enrichment Analysis (ssGSEA) (escape version 1.12.0). Significant Hallmark gene sets were defined as those with an FDR < 0.1.

## Re-analysis of bulk RNA sequencing

We also include publicly available bulk RNA-seq data (GEO: GSE196311, GSE130807)^2525^. Raw FASTQ files were processed using Trim Galore (v0.6.10) and STAR (v2.7.10a). Read counts were generated using featureCounts (Subread v2.0.1). The R package DESeq2 (v1.38.3), and iDEP web tool (v2.01)^69^ were used to normalize the gene expression matrix. Gene Set Enrichment Analysis (GSEA) was performed using GSEA software (v4.3.3), focusing on Hallmark gene sets from the MSigDB collection. Significant Hallmark gene sets were defined as those with an FDR < 0.1.

## Re-analysis of single-cell RNA sequencing data from *Mesp1^+^* CPM cells

We re-analyzed scRNA-seq data from *Mesp1^+^* CPM cells between E8.0 and E10.5 (GEO: GSE167493)^37^. Raw reads were processed using 10x Genomics Cell Ranger software (v6.1.2), mapped to GRCm38 (mm10), and feature-barcode matrices were generated on the SHIROKANE SC. Data analysis was conducted using R (v4.2.0) and the Seurat package (v4.3.0). Quality control yielded the following cell counts: E8.0 (8,276 cells), E8.25 (3,255 cells), E9.5 (4,288 cells), and E10.5 (9,664 cells). Cells with 1,000–7,500 feature counts and < 5% mitochondrial genes were retained, leaving 22,899 cells for analysis. Normalization was performed using *sctransform*, selecting the top 3,000 variable genes. Cell cycle effects were regressed using phase scores based on markers from Tirosh et al. (2015, updated in 2019)^68^. Two approaches were used: one across all stages (E8.0–E10.5) and one focusing on early stages (E8.0–E8.25). Batch correction and integration were conducted using Anchor-based CCA, followed by dimensionality reduction employing PCA and clustering with k-nearest neighbor (k-NN) graphs (E8.0–E10.5: 20 clusters, E8.0–E8.25: 15 clusters). Clustering was visualized using UMAP. Clusters were annotated based on marker genes identified using Seurat’s *FindAllMarkers* function (FDR < 0.05, log2-fold change > 1). Subclustering of EC clusters was performed in both the E8.0–E10.5 (C7 and 15) and E8.0–E8.25 (C3, 4, 11, and 12) approaches, following the outline reported by Piper et al^70^. ECs were extracted using Seurat’s *subset* function, and the downstream analysis (PCA, clustering, and visualization) was performed. The E8.0–E10.5 approach identified eight clusters (C0–7), while the E8.0–E8.25 approach identified seven clusters (C0–6). The following parameters were used: for E8.0– E10.5, PCs 1 to 20, k = 20, resolution = 0.6; for E8.0–E8.25, PCs 1 to 5, k = 20, resolution = 0.5. Clusters were visualized using UMAP and annotated based on marker genes identified using Seurat’s *FindAllMarkers* (FDR < 0.05, log2-fold change > 1).

RNA velocity analysis was performed following the velocyto and scVelo tutorials^71,72^. We used velocyto (v0.17.17) to separate spliced and unspliced mRNA reads from the scRNA-seq mapping data, generating spliced and unspliced count matrices on the SHIROKANE SC.

Subsequent RNA velocity estimation, pseudotime calculation, and trajectory inference were performed using scVelo (v0.3.3) in Python (v3.11.7), based on the spliced/unspliced matrices and Seurat analysis results. The following parameters were used for both the E8.0–E10.5 and E8.0–E8.25 approaches: PCs 1 to 30, k = 30 NN graph. Pseudotime was calculated from the directed velocity graph, and the inferred differentiation trajectories were represented as a PAGA graph, extended with velocity-derived directionality.

## Statistics and reproducibility

GraphPad Prism 10 was used for graphical representation and statistical analysis. Data are presented as the mean ± standard error of the mean (SEM). Comparisons between two groups were made using Mann–Whitney U tests. P-values less than 0.05 were considered significant. The experiments were not randomized, and blinding was not performed during analysis or quantification. No statistical methods were used to predetermine sample sizes.

## Data availability

Data availability for scRNA-seq data, raw data will be available in GEO (GSE 279129). There is no restriction on data availability. Source data are provided in this paper.

## AUTHORS’ CONTRIBUTIONS

K.M. and K.I.-Y. conceived the study and designed the experiments. S.T., N.N., and K.M. performed the experiments. Single-cell RNA sequencing analysis was conducted by S.T. and reviewed by K.N. and K.M. K. N. confirmed some of the data, obtaining similar results. Y.K. provided the *CDH5-CreERT2* mice. O.N. supplied the *Mef2c-AHF-Cre* mice. Y.H., K.H., S.T., and E.M. provided human samples. The pathological diagnosis of vascular malformations was conducted by Y.H. and K.H. H.U. and M.N., specialists in vascular malformation surgical treatment, reviewed the manuscript from a clinical perspective. K.M. coordinated the experimental work, analyzed the data, and wrote the manuscript with contributions from all authors.

## ACKNOWLEDGMENTS

We extend our sincere gratitude to all laboratory members for their insightful discussions and encouragement throughout this study. This work was supported in part by Grants-in-Aid for Scientific Research from the Ministry of Education, Culture, Sports, Science, and Technology of Japan (20K17072 and 23K15949 to K.M.); the Japan Foundation for Applied Enzymology (VBIC to K.M.); the SENSHIN Medical Research Foundation (K.M.); the Takeda Science Foundation (K.M.); The Ichiro Kanehara Foundation for the Promotion of Medical Sciences and Medical Care (K.M.); the Mie University Life Relay Foundation (K.M.); the HITACHI Group Foundation (Kurata Research Grant to K.M.); the Japan Intractable Diseases Research Foundation (K.M.); the Mochida Memorial Foundation for Medical and Pharmaceutical Research (K.M.); the Japan Agency for Medical Research and Development (AMED) under Grant Number 22jm0610079h0001 (K.M.); the Mie University Research Promotion and Graduate School Reform-Related Research Grant Project (K.M.); and the TERUMO Life Science Foundation (K.M.); The Sumitomo Foundation (K.M.). We also acknowledge the NGS Core Facility at the Research Institute for Microbial Diseases of Osaka University for their support with sequencing.

---

**Supplemental Table 1:**
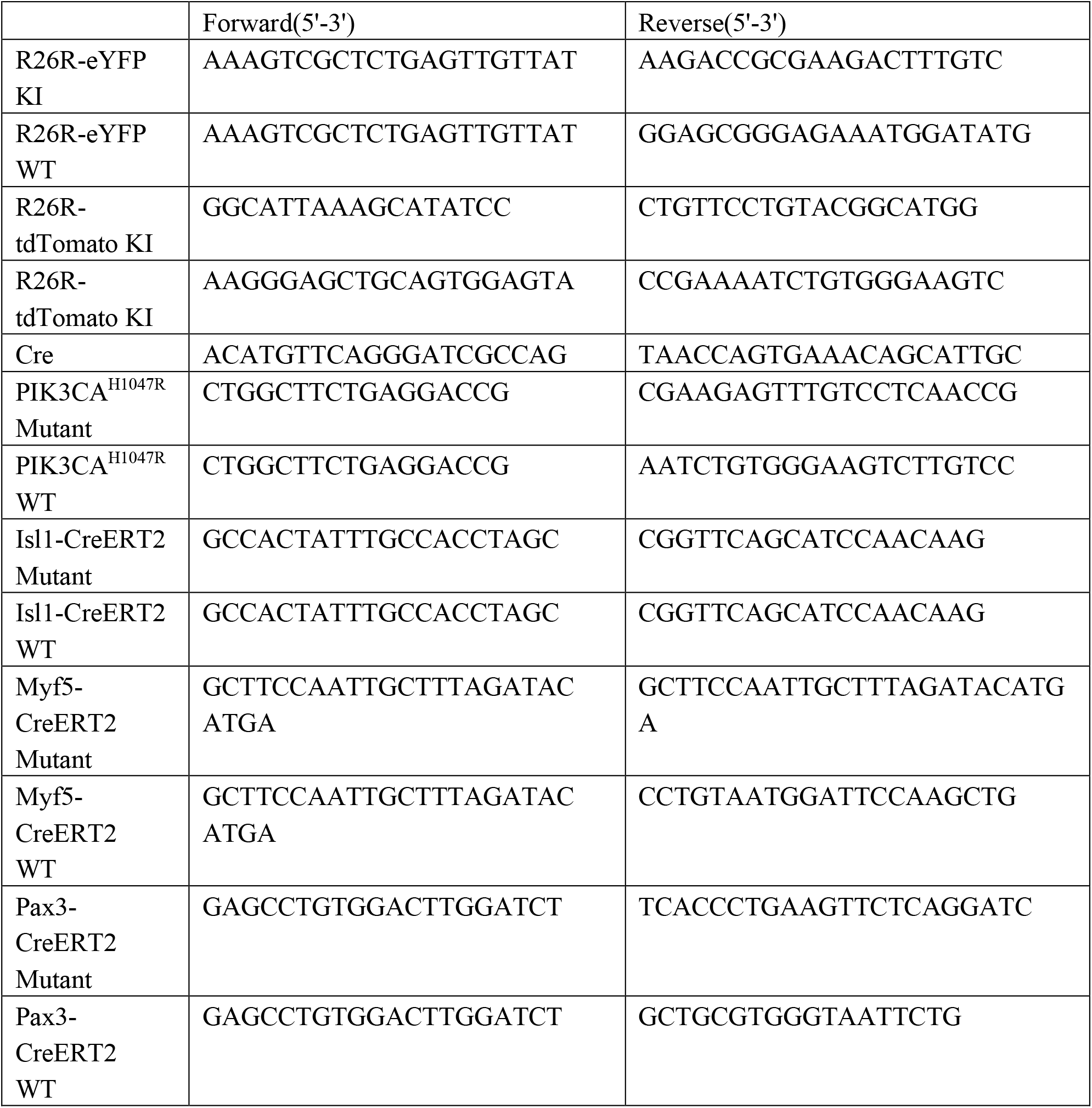
Primers used for genotyping

## References

1. Pang, C., Lim, C. S., Brookes, J., Tsui, J. & Hamilton, G. Emerging importance of molecular pathogenesis of vascular malformations in clinical practice and classifications. Vasc. Med. 25, 364–377 (2020).

2. Castillo, S. D., Baselga, E. & Graupera, M. PIK3CA mutations in vascular malformations. Curr. Opin. Hematol. 26, 170–178 (2019).

3. Limaye, N., Boon, L. M. & Vikkula, M. From germline towards somatic mutations in the pathophysiology of vascular anomalies. Hum. Mol. Genet. 18, R65–R74 (2009).

4. Zenner, K. et al. Genotype correlates with clinical severity in PIK3CA-associated lymphatic malformations. Jci Insight 4, (2019).

5. Lee, J. W. & Chung, H. Y. Vascular anomalies of the head and neck: current overview. Arch. Craniofacial Surg. 19, 243–247 (2018).

6. Castillo, S. D., Baselga, E. & Graupera, M. PIK3CA mutations in vascular malformations. Curr Opin Hematol 26, 170–178 (2019).

7. Damme, A. V., Seront, E., Dekeuleneer, V., Boon, L. M. & Vikkula, M. New and Emerging Targeted Therapies for Vascular Malformations. Am. J. Clin. Dermatol. 21, 657– 668 (2020).

8. Mimura, H. et al. Japanese clinical practice guidelines for vascular anomalies 2017. Pediatr Int 62, 260–307 (2020).

9. Kobialka, P. et al. The onset of PI3K-related vascular malformations occurs during angiogenesis and is prevented by the AKT inhibitor miransertib. Embo Mol Med 14, e15619 (2022).

10. Boscolo, E. et al. AKT hyper-phosphorylation associated with PI3K mutations in lymphatic endothelial cells from a patient with lymphatic malformation. Angiogenesis 18, 151–162 (2015).

11. Luks, V. L. et al. Lymphatic and Other Vascular Malformative/Overgrowth Disorders Are Caused by Somatic Mutations in PIK3CA. J Pediatrics 166, 1048–1054.e5 (2015).

12. Mäkinen, T., Boon, L. M., Vikkula, M. & Alitalo, K. Lymphatic Malformations: Genetics, Mechanisms and Therapeutic Strategies. Circ Res 129, 136–154 (2021).

13. Limaye, N. et al. Somatic Activating PIK3CA Mutations Cause Venous Malformation. Am. J. Hum. Genet. 97, 914–921 (2015).

14. Hirose, K. et al. Comprehensive phenotypic and genomic characterization of venous malformations. Hum. Pathol. (2024) doi:10.1016/j.humpath.2024.02.004.

15. Castel, P. et al. Somatic PIK3CA mutations as a driver of sporadic venous malformations. Sci Transl Med 8, 332ra42 (2016).

16. Castillo, S. D. et al. Somatic activating mutations in Pik3ca cause sporadic venous malformations in mice and humans. Sci Transl Med 8, 332ra43 (2016).

17. Graupera, M. et al. Angiogenesis selectively requires the p110α isoform of PI3K to control endothelial cell migration. Nature 453, 662–666 (2008).

18. Gupta, S. et al. Binding of Ras to Phosphoinositide 3-Kinase p110α Is Required for Ras- Driven Tumorigenesis in Mice. Cell 129, 957–968 (2007).

19. Stanczuk, L. et al. cKit Lineage Hemogenic Endothelium-Derived Cells Contribute to Mesenteric Lymphatic Vessels. Cell Reports 10, 1708–1721 (2015).

20. Blasio, L. di et al. PI3K/mTOR inhibition promotes the regression of experimental vascular malformations driven by PIK3CA-activating mutations. Cell Death Dis 9, 45 (2018).

21. Madsen, R. R., Vanhaesebroeck, B. & Semple, R. K. Cancer-Associated PIK3CA Mutations in Overgrowth Disorders. Trends Mol Med 24, 856–870 (2018).

22. Keppler-Noreuil, K. M., et al. PIK3CA-related overgrowth spectrum (PROS): Diagnostic and testing eligibility criteria, differential diagnosis, and evaluation. Am. J. Méd. Genet. Part A 167, 287–295 (2015).

23. Gomes, I. P. et al. Assessment of PI3K/AKT and MAPK/ERK pathways activation in oral lymphatic malformations. *Oral Surg., Oral Med., Oral Pathol*. Oral Radiol. 133, 216– 220 (2022).

24. Blesinger, H. et al. PIK3CA mutations are specifically localized to lymphatic endothelial cells of lymphatic malformations. Plos One 13, e0200343 (2018).

25. Jauhiainen, S. et al. ErbB signaling is a potential therapeutic target for vascular lesions with fibrous component. eLife 12, e82543 (2023).

26. Aw, W. Y. et al. Microphysiological model of PIK3CA-driven vascular malformations reveals a role of dysregulated Rac1 and mTORC1/2 in lesion formation. Sci. Adv. 9, eade8939 (2023).

27. Broek, R. W. T. et al. Comprehensive molecular and clinicopathological analysis of vascular malformations: A study of 319 cases. *Genes, Chromosom*. Cancer 58, 541–550 (2019).

28. Seront, E., Damme, A. V., Boon, L. M. & Vikkula, M. Rapamycin and treatment of venous malformations. Curr. Opin. Hematol. 26, 185–192 (2019).

29. Rodriguez-Laguna, L. et al. Somatic activating mutations in PIK3CA cause generalized lymphatic anomalyPIK3CA mutations in generalized lymphatic anomaly. J Exp Medicine 216, 407–418 (2019).

30. Boscolo, E. et al. Rapamycin improves TIE2-mutated venous malformation in murine model and human subjects. J. Clin. Investig. 125, 3491–3504 (2015).

31. Maruani, A. et al. Sirolimus (Rapamycin) for Slow-Flow Malformations in Children. JAMA Dermatol. 157, 1289–1298 (2021).

32. Maruyama, K., Miyagawa-Tomita, S., Mizukami, K., Matsuzaki, F. & Kurihara, H. Isl1- expressing non-venous cell lineage contributes to cardiac lymphatic vessel development. Dev Biol 452, 134–143 (2019).

33. Maruyama, K. et al. The cardiopharyngeal mesoderm contributes to lymphatic vessel development in mouse. Elife 11, (2022).

34. Yamaguchi, S. et al. The development of early human lymphatic vessels as characterized by lymphatic endothelial markers. EMBO J. 1–18 (2024) doi:10.1038/s44318-024-00045-0.

35. Lioux, G. et al. A Second Heart Field-Derived Vasculogenic Niche Contributes to Cardiac Lymphatics. Dev Cell 52, 350–363 (2020).

36. Lupu, I.-E. et al. Direct specification of lymphatic endothelium from non-venous angioblasts. Biorxiv 2022.05.11.491403 (2022) doi:10.1101/2022.05.11.491403.

37. Nomaru, H. et al. Single cell multi-omic analysis identifies a Tbx1-dependent multilineage primed population in murine cardiopharyngeal mesoderm. Nat Commun 12, 6645 (2021).

38. Zhang, M. et al. Coronary vessels contribute to de novo endocardial cells in the endocardium-depleted heart. Cell Discov. 9, 4 (2023).

39. Yu, Y. et al. Targeting AKT1-E17K and the PI3K/AKT Pathway with an Allosteric AKT Inhibitor, ARQ 092. PLoS ONE 10, e0140479 (2015).

40. Mirzaa, G., et al. PIK3CA-associated developmental disorders exhibit distinct classes of mutations with variable expression and tissue distribution. JCI Insight 1, e87623 (2016).

41. Brouillard, P. et al. Non-hotspot PIK3CA mutations are more frequent in CLOVES than in common or combined lymphatic malformations. Orphanet J. Rare Dis. 16, 267 (2021).

42. Roy, A. et al. Mouse models of human PIK3CA-related brain overgrowth have acutely treatable epilepsy. eLife 4, e12703 (2015).

43. Peyre, M. et al. Somatic PIK3CA Mutations in Sporadic Cerebral Cavernous Malformations. N. Engl. J. Med. 385, 996–1004 (2021).

44. Ren, A. A. et al. PIK3CA and CCM mutations fuel cavernomas through a cancer-like mechanism. Nature 594, 271–276 (2021).

45. Orsenigo, F. et al. Mapping endothelial-cell diversity in cerebral cavernous malformations at single-cell resolution. eLife 9, e61413 (2020).

46. Luo, W. et al. Arterialization requires the timely suppression of cell growth. Nature 589, 437–441 (2021).

47. Hong, C. C., Peterson, Q. P., Hong, J.-Y. & Peterson, R. T. Artery/Vein Specification Is Governed by Opposing Phosphatidylinositol-3 Kinase and MAP Kinase/ERK Signaling. Curr. Biol. 16, 1366–1372 (2006).

48. Zhong, H. et al. Modulation of hypoxia-inducible factor 1alpha expression by the epidermal growth factor/phosphatidylinositol 3-kinase/PTEN/AKT/FRAP pathway in human prostate cancer cells: implications for tumor angiogenesis and therapeutics. Cancer Res. 60, 1541–5 (2000).

49. Xiao, Y. et al. PDGF Promotes the Warburg Effect in Pulmonary Arterial Smooth Muscle Cells via Activation of the PI3K/AKT/mTOR/HIF-1α Signaling Pathway. Cell. Physiol. Biochem. 42, 1603–1613 (2017).

50. Fitzgerald, G., Soro-Arnaiz, I. & Bock, K. D. The Warburg Effect in Endothelial Cells and its Potential as an Anti-angiogenic Target in Cancer. Front. Cell Dev. Biol. 6, 100 (2018).

51. Yu, P. et al. FGF-dependent metabolic control of vascular development. Nature 545, 224–228 (2017).

52. Zou, Z., Tao, T., Li, H. & Zhu, X. mTOR signaling pathway and mTOR inhibitors in cancer: progress and challenges. Cell Biosci. 10, 31 (2020).

53. ISSVA classification for vascular anomalies ©.

54. Hori, Y. et al. PIK3CA mutation correlates with mTOR pathway expression but not clinical and pathological features in Fibro-adipose vascular anomaly (FAVA). Diagn Pathol 17, 19 (2022).

55. Kisanuki, Y. Y. et al. Tie2-Cre Transgenic Mice: A New Model for Endothelial Cell- Lineage Analysis in Vivo. Dev Biol 230, 230–242 (2001).

56. Okabe, K. et al. Neurons Limit Angiogenesis by Titrating VEGF in Retina. Cell 159, 584–596 (2014).

57. Cai, C.-L. et al. Isl1 Identifies a Cardiac Progenitor Population that Proliferates Prior to Differentiation and Contributes a Majority of Cells to the Heart. Dev Cell 5, 877–889 (2003).

58. Verzi, M. P., McCulley, D. J., Val, S. D., Dodou, E. & Black, B. L. The right ventricle, outflow tract, and ventricular septum comprise a restricted expression domain within the secondary/anterior heart field. Dev Biol 287, 134–145 (2005).

59. Laugwitz, K.-L. et al. Postnatal isl1+ cardioblasts enter fully differentiated cardiomyocyte lineages. Nature 433, 647–653 (2005).

60. Biressi, S. et al. Myf5 expression during fetal myogenesis defines the developmental progenitors of adult satellite cells. Dev Biol 379, 195–207 (2013).

61. Southard, S. et al. A series of Cre-ERT2 drivers for manipulation of the skeletal muscle lineage. *genesis* **52**, 759–770 (2014).

62. Srinivas, S. et al. Cre reporter strains produced by targeted insertion of EYFP and ECFP into the ROSA26 locus. Bmc Dev Biol 1, 4 (2001).

63. Madisen, L. et al. A robust and high-throughput Cre reporting and characterization system for the whole mouse brain. Nat Neurosci 13, 133–140 (2010).

64. Adams, J. R. et al. Cooperation between Pik3ca and p53 Mutations in Mouse Mammary Tumor Formation. Cancer Res 71, 2706–2717 (2011).

65. Lee, J.-Y. et al. Pharmacokinetic Characterization of LW6, a Novel Hypoxia-Inducible Factor-1α (HIF-1α) Inhibitor in Mice. Molecules 26, 2226 (2021).

66. Wu, F., Tamhane, M. & Morris, M. E. Pharmacokinetics, Lymph Node Uptake, and Mechanistic PK Model of Near-Infrared Dye-Labeled Bevacizumab After IV and SC Administration in Mice. AAPS J. 14, 252–261 (2012).

67. Martinez-Corral, I. et al. Blockade of VEGF-C signaling inhibits lymphatic malformations driven by oncogenic PIK3CA mutation. Nat Commun 11, 2869 (2020).

68. Kowalczyk, M. S. et al. Single-cell RNA-seq reveals changes in cell cycle and differentiation programs upon aging of hematopoietic stem cells. Genome Res. 25, 1860– 1872 (2015).

69. Ge, S. X., Son, E. W. & Yao, R. iDEP: an integrated web application for differential expression and pathway analysis of RNA-Seq data. Bmc Bioinformatics 19, 534 (2018).

70. Piper, R. J. et al. Towards network-guided neuromodulation for epilepsy. Brain 145, 3347–3362 (2022).

71. Manno, G. L. et al. RNA velocity of single cells. Nature 560, 494–498 (2018).

72. Bergen, V., Lange, M., Peidli, S., Wolf, F. A. & Theis, F. J. Generalizing RNA velocity to transient cell states through dynamical modeling. Nat. Biotechnol. 38, 1408–1414 (2020).

