## Supplemental Figure and legends for "Embryological cellular origins and hypoxia-mediated mechanisms in *PIK3CA*-Driven refractory vascular malformations"

**Supplemental Figure 1**

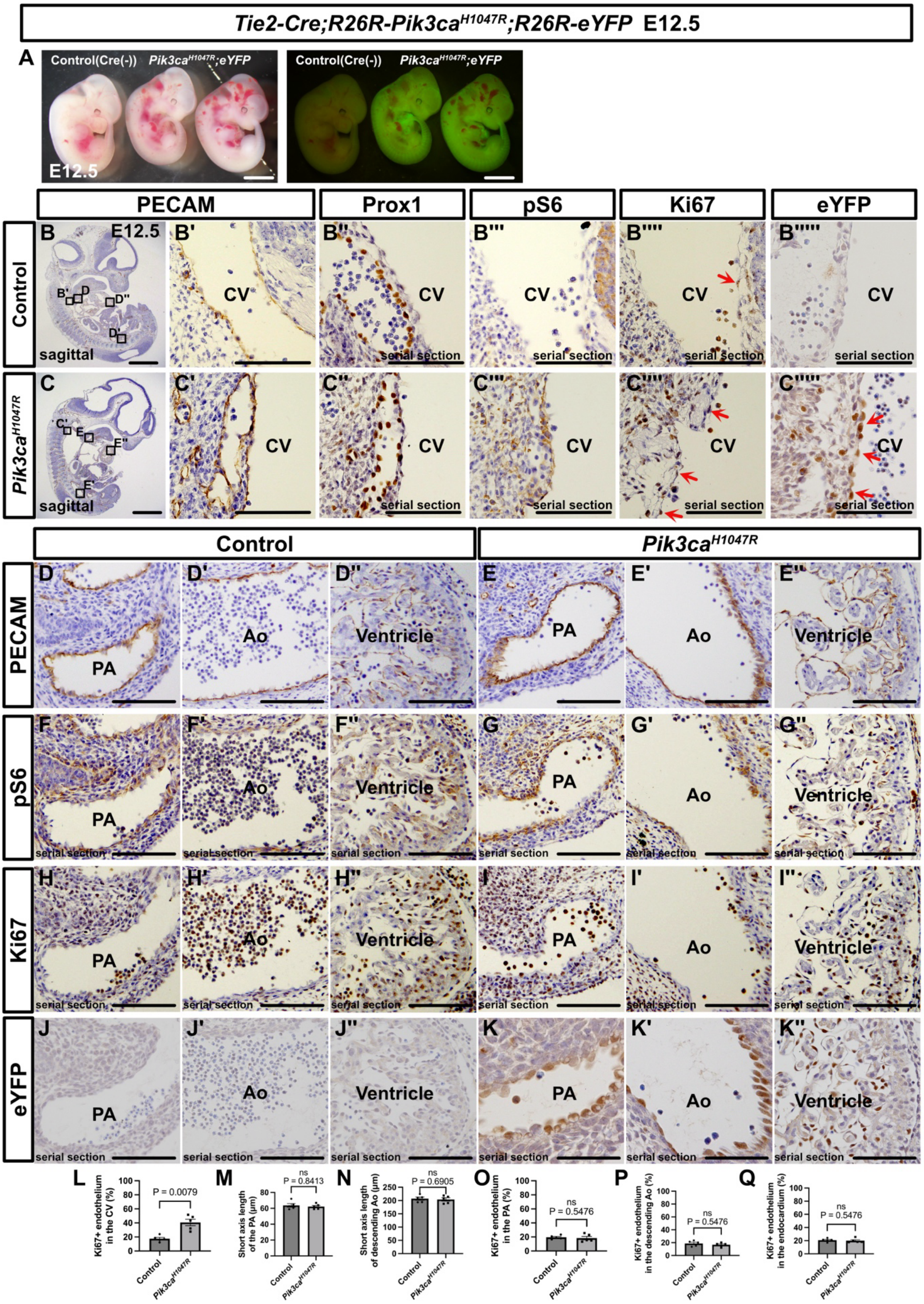

**Supplemental Figure 1. Pik3ca$^{H1047R}$ expression in *Tie2-Cre* embryos does not significantly affect the endothelium of the aorta, pulmonary artery, or endocardium.**

**(A)** Gross morphology of the control and *Tie2-Cre; R26R-Pik3ca$^{H1047R}$* mutant embryos at E12.5. In the control, eYFP is not expressed, but in the mutant, eYFP is expressed throughout the body. **(B-K”)** Immunostaining of sagittal sections with the indicated antibodies. An increase in Ki67$^{+}$ cells in the cardinal vein ECs is observed (**B”””, C”””**, red arrows). In mutant mice, eYFP$^{+}$ cells are observed in both cardinal vein ECs and LECs (**B””’, C””’**, red arrows). **(D-K”)** Similar analysis in the pulmonary artery, descending aorta, and endocardium. CV, cardinal vein; Ao, aorta; PA, pulmonary artery. Each dot represents a value obtained from one sample. Scale bars, 100 μm **(B’-B””’, C’-E””’, D-K”)** and 1 mm **(A)**. The nonparametric Mann–Whitney U test was used for statistical analysis, with exact p-values indicated. ns ≥ 0.05.

### Supplemental Figure 2

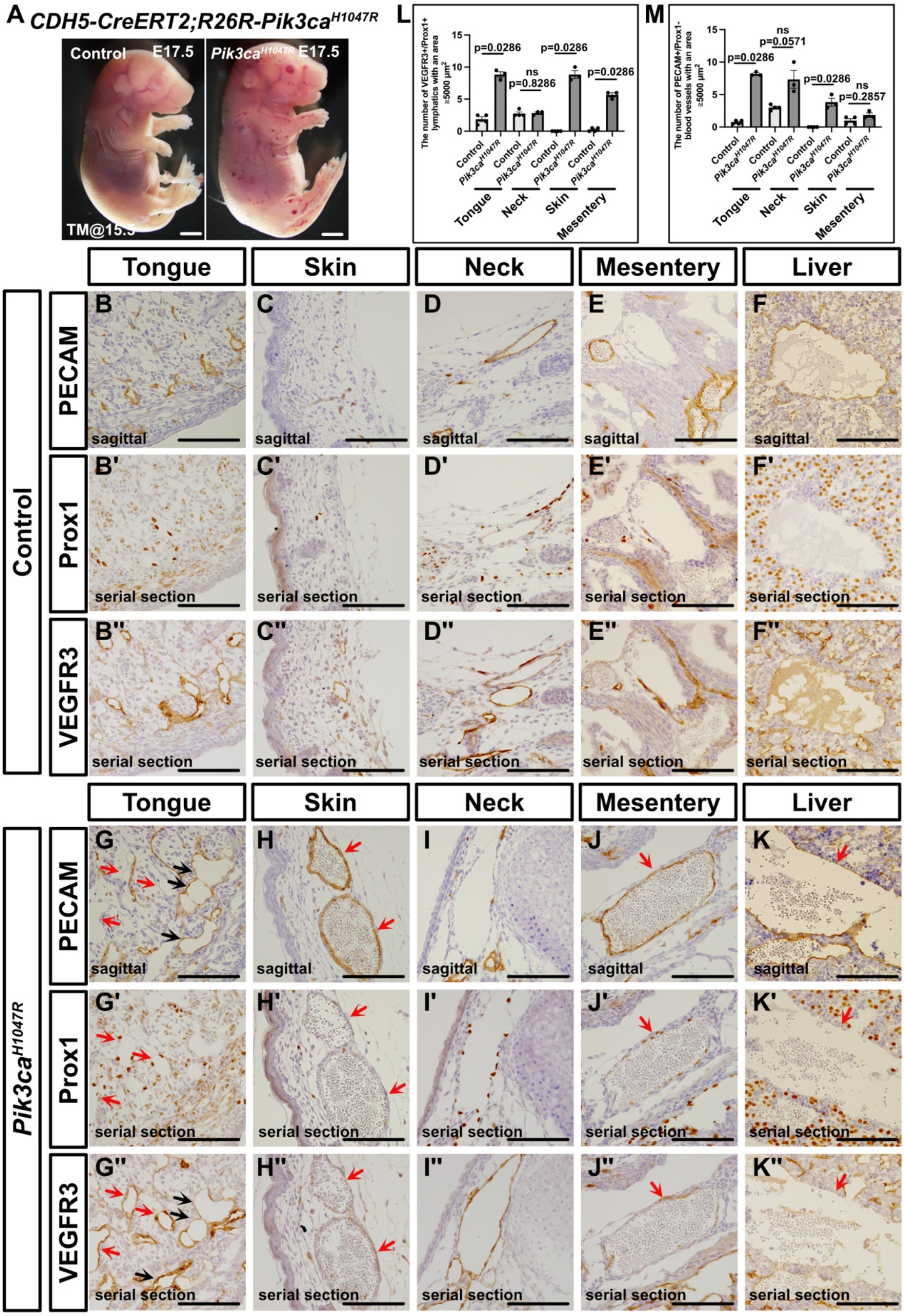

**Supplemental Figure 2. Pik3caH1047R expression in endothelial cells at E15.5 leads to systemic vascular malformations.**

**(A)** Gross morphology of the control and *CDH5-CreERT2; R26R-Pik3ca*$^{H1047R}$ mutant embryos at E17.5, with tamoxifen administered to pregnant mice at E15.5. **(B-K")** Sagittal section immunohistochemistry using the indicated antibodies. **(B-B", G-G")** In mutant embryos, mildly dilated $PECAM^-$/$Prox1^+$/$VEGFR3^+$ lymphatic vessels (red arrows) and an increased number of $PECAM^+$/$Prox1^-$/$VEGFR3^+$ blood vessels (black arrows) are observed in the tongue. **(C-C", H-H")** Aberrant, dilated $PECAM^+$/partially $Prox1^+$/partially $VEGFR3^+$ vessels are seen in the skin (red arrows). **(D-D", I-I")** No significant differences in the neck (larynx) are observed between the control and mutant embryos. **(E-E", J-J")** In the mesentery, aberrant, dilated $PECAM^+$/partially $Prox1^+$/partially $VEGFR3^+$ vessels similar to those in the skin are seen (red arrows). **(F-F", K-K")** In the liver, dilated $PECAM^+$/$Prox1^-$/partially $VEGFR3^+$ blood vessels are observed. The number of PECAM+ vessels with an area ≥ 20,000 $\mu m^2$ in the liver is 0 ± 0 (median ± SEM) (n = 5) in controls and 1.5 ± 0.33 (median ± SEM) (n = 3) in mutants (p = 0.0179). **(L, M)** Statistical analysis in the tongue, neck, skin, and mesentery. Each dot represents a value obtained from one sample. Scale bars: 100 µm **(B-K")** and 2 mm **(A)**. The nonparametric Mann–Whitney U test was used for statistical analysis. ns ≥ 0.05.

### Supplemental Figure 3

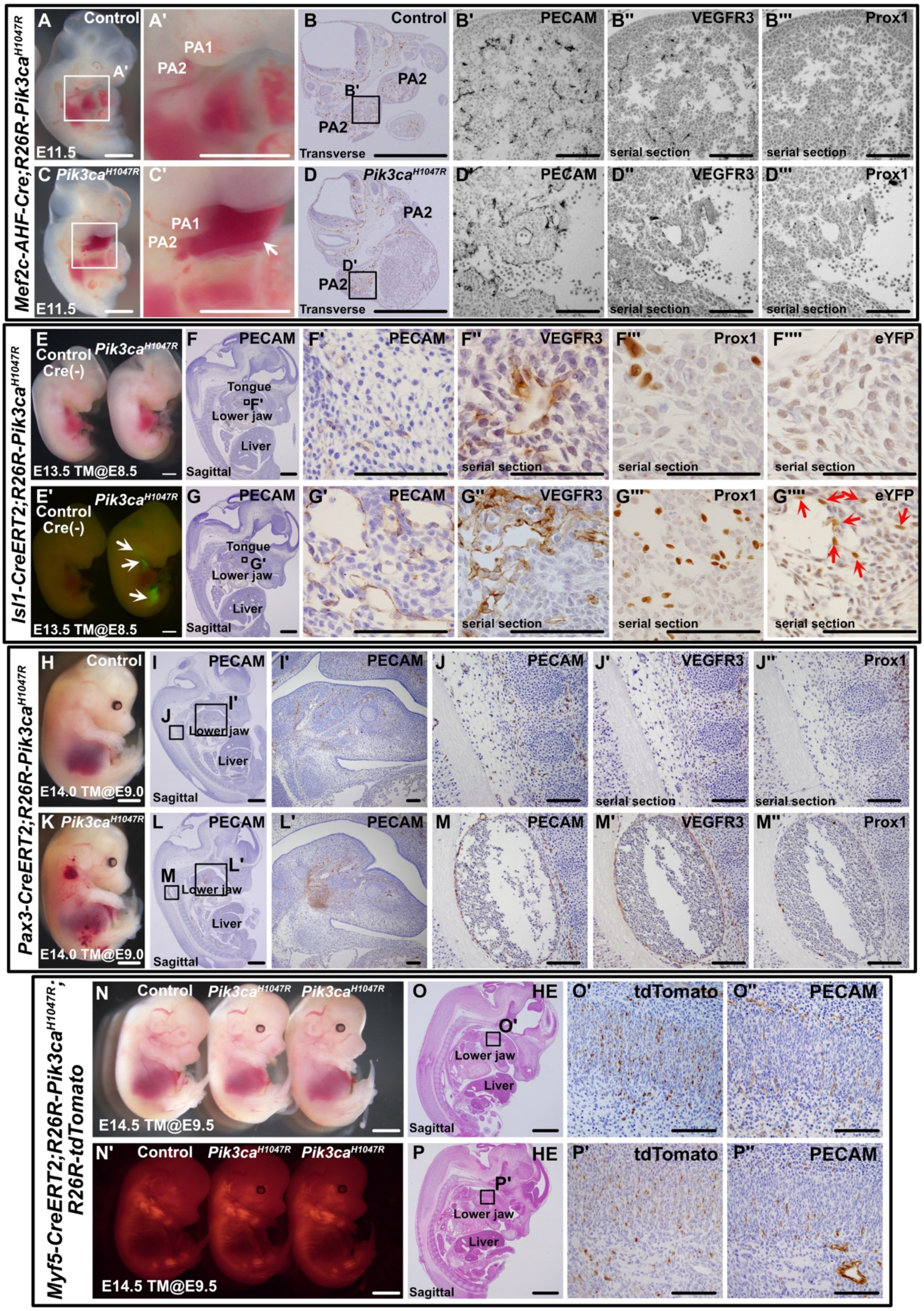

**Supplemental Figure 3. Pik3ca$^{H1047R}$ expression in CPM induces vascular malformations in the head and neck.**

**(A, A’, C, C’)** Gross morphology of the control and *Mef2c-AHF-Cre; R26R-Pik3ca*$^{H1047R}$ mutant embryos at E11.5. **(B-B”’, D-D”’)** Transverse section immunostaining with the indicated antibodies. Mutant embryos show dilated PECAM$^+$/Prox1$^-$/partially VEGFR3$^+$ vessels extending from the first to the second pharyngeal arches (control vs. mutant PECAM$^+$ vasculature area in PA2 at E11.5 (μm$^2$): 4203 ± 480.88 (median ± SEM) (n = 4) vs. 335176 ± 102003.10 (n = 4), p = 0.0286). **(E, E’)** Gross morphology of the control and *Isl1-CreERT2; R26R-Pik3ca*$^{H1047R}$*; R26R-eYFP* mutant embryos at E13.5, after tamoxifen administration at E8.5. eYFP expression is observed from the lower jaw to the neck and outflow tracts, and in the genital region (white arrows). **(F-F””, G-G””)** Mutant embryos show dilated PECAM$^+$/Prox1$^+$/VEGFR3$^+$ lymphatic vessels between the lower jaw and tongue (control vs. mutant Prox1$^+$ lymphatic vessel area in the lower jaw and tongue at E13.5 (μm$^2$): 2296 ± 167.26 (median ± SEM) (n = 5) vs. 17962 ± 851.35 (n = 5), p = 0.0079)**.** These vessels are also eYFP$^+$ (red arrows). **(H, K)** Gross morphology of the control and *Pax3-CreERT2; R26R-Pik3ca*$^{H1047R}$ mutant embryos at E14.0, after tamoxifen administration at E9.0. **(I-M”)** No vascular malformations are seen in the head and neck of mutant embryos, but dilated PECAM$^+$/partially Prox1$^+$/VEGFR3$^+$ blood-filled vessels are observed around the spine. (control vs. mutant PECAM+ vessels with an area ≥ 5000 μm$^2$ around the spine: 0 ± 0 (median ± SEM) (n = 5) vs. 2 ± 0.187 (median ± SEM) (n = 5), p = 0.0079). **(N, N’)** Gross morphology of the control and *Myf5-CreERT2; R26R-Pik3ca*$^{H1047R}$*; R26R-tdTomato* mutant embryos at E14.5, after tamoxifen administration at E9.5. **(O, P)** Sagittal section H&E staining. **(O’, O”, P’, P”)** Sagittal section immunostaining with the indicated antibodies. Scale bars: 100 μm **(B’-B”’, D’-D”’, F’-F””, G’-G””, I’-J”, L’-M”, O’-P”)**, 1 mm **(A-B, C-D, F, G, H, I, K, L, O, P)**, and 2 mm **(E, E’, N, N’)**. The nonparametric Mann–Whitney U test was used for statistical analysis.

69 **Supplemental Figure 4**

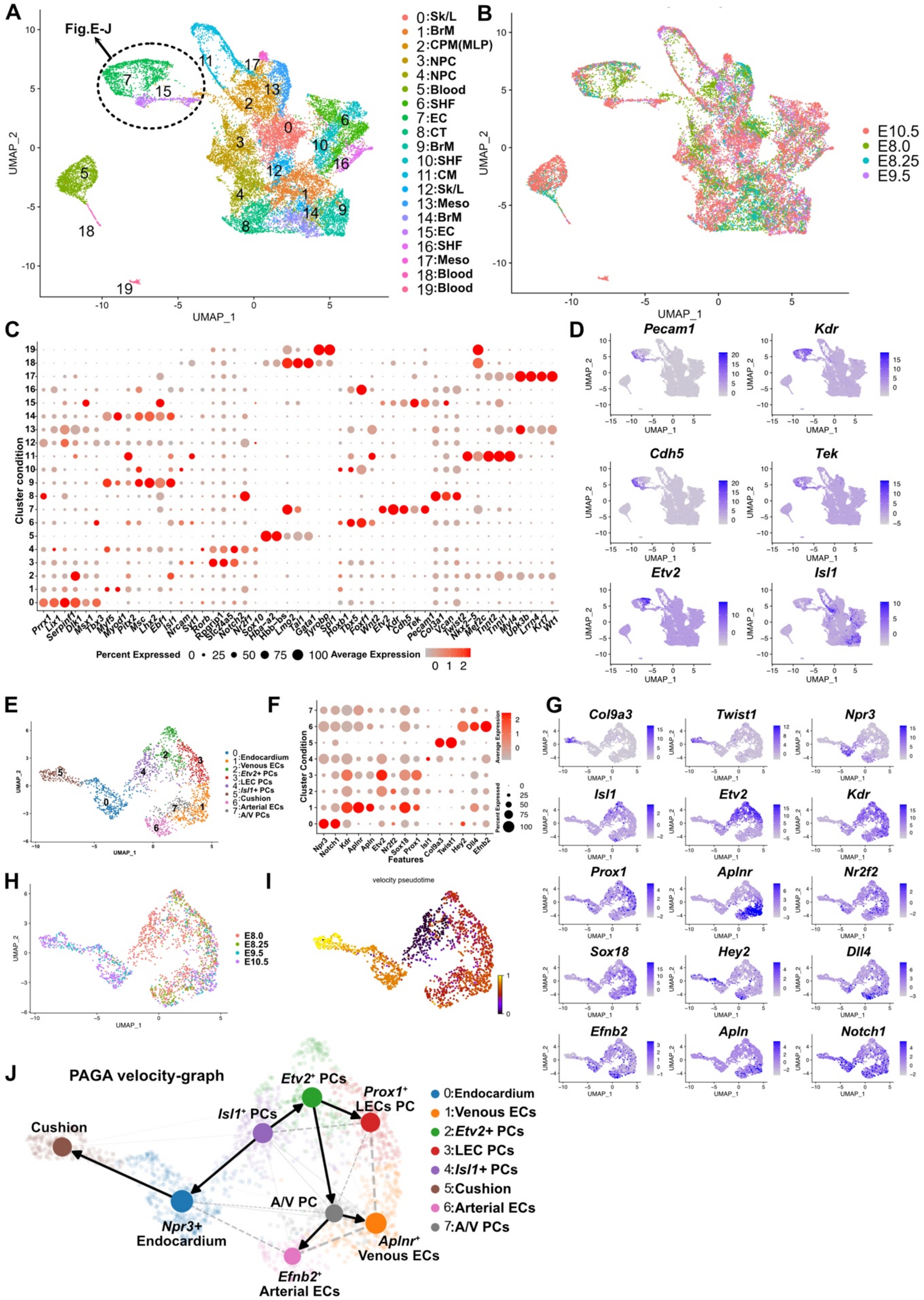


70

**Supplemental Figure 4. Predicted differentiation lineage of endothelial cells from the CPM.**

**(A-G)** Re-analysis of scRNA-seq data[37] of *Mesp1*$^{+}$ CPM at E8.0-E10.5. (A) UMAP plot with clusters color-coded (0-19): 0: Sk/L, skeleton/limb progenitor cells; 1: BrM, branchial muscle; 2: CPM (MLP), cardiopharyngeal mesoderm (multilineage progenitor cells); 3, 4: NPC, neural progenitor cells; 5: blood, blood cells; 6: SHF, second heart field; 7: EC, endothelial cells; 8: CT, Connective tissue; 9: BrM, branchial muscle; 10: SHF, second heart field; 11: CM, cardiomyocyte; 12: Sk/L, skeleton/limb progenitor cells; 13: Meso, mesothelium; 14: BrM, branchial muscle; 15: EC, endothelial cell; 16: SHF, second heart field; 17: Meso, mesothelium; 18,19: blood, blood cells. Black dotted circle indicates endothelial clusters, which underwent sub-clustering analysis. **(B)** UMAP plot color-coded by embryonic day: green (E8.0), light blue (E8.25), purple (E9.5), and red (E10.5). **(C)** Heatmap displaying the average expression levels of marker genes per cluster. **(D)** UMAP plot showing the expression levels of endothelial markers. (**E**) UMAP plot with clusters color-coded (0-7): 0: Endocardium; 1: Venous EC, venous endothelial cells; 2: *Etv2*$^{+}$ PCs, *Etv2*$^{+}$ endothelial progenitor cells; 3: LEC PCs, lymphatic endothelial progenitor cells; 4: *Isl1*$^{+}$ PCs, *Isl1*$^{+}$ endothelial progenitor cells; 5: Cushion, cushion tissue; 6: Arterial ECs, arterial endothelial cells; 7: A/V PCs, arterial and venous endothelial progenitor cells. (**F**) Heatmap displaying the average expression levels of marker genes per cluster. (**G**) Heatmap displaying the average expression levels of marker genes per cluster. (**H**) UMAP plot, color-coded by embryonic day, with red (E8.0), green (E8.25), light blue (E9.5), and purple (E10.5). **(I)** UMAP plot representing pseudotime calculated from RNA velocity analysis. (**J**) PAGA graph illustrating the predicted differentiation lineage of CPM-derived endothelial cells based on RNA velocity.

95 **Supplemental Figure 5**

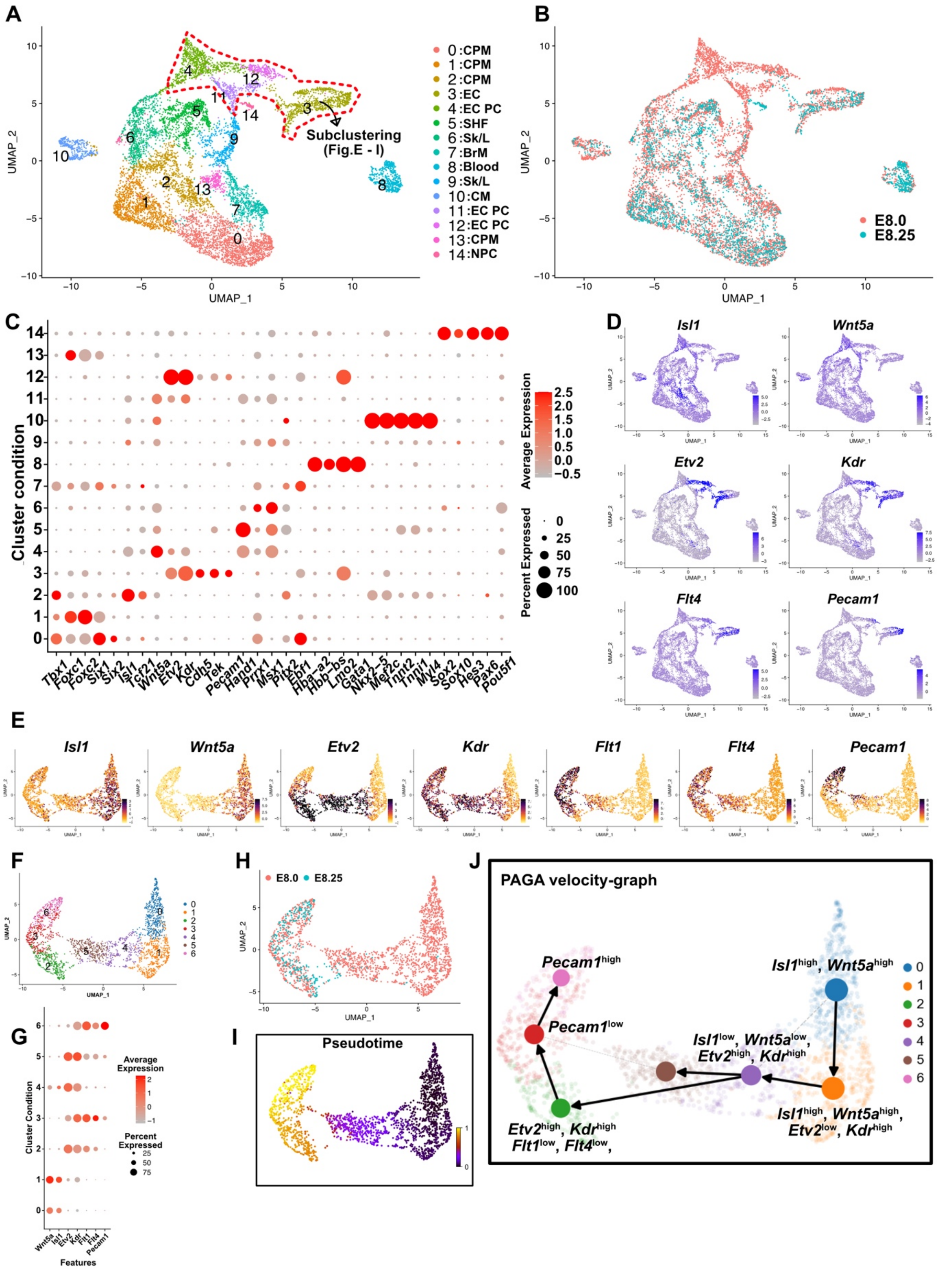


96

**Supplemental Figure 5. The downregulation of *Isl1* and the expression of *Etv2* drive the differentiation of endothelial cells from the CPM.**

(A) UMAP plot color-coded by cluster (0-14): 0, 1, 2: CPM, cardiopharyngeal mesoderm; 3: EC, endothelial cells; 4: EC PC, endothelial cell progenitors; 5: SHF, second heart field; 6, 9: Sk/L, skeleton/limb progenitor cells; 7: BrM, branchial muscle; 8: blood, blood cell; 9: BrM, branchial muscle; 10: CM, cardiomyocyte; 11, 12: EC PC, endothelial cell progenitors; 13: CPM, cardiopharyngeal mesoderm; 14: NPC, neural progenitor cells. (B) UMAP plot color-coded by embryonic day, with red representing E8.0 and light blue representing E8.25. (C) Heatmap showing the average expression levels of marker genes for each cluster. (D) UMAP plot displaying the expression levels of CPM or endothelial cell marker genes. (E) Sub-clustering analysis of clusters 3, 4, 11, and 12, showing the endothelial cell marker genes. (F) UMAP plot with clusters color-coded (0-6). (G) Heatmap showing the average expression levels of marker genes for each cluster. (H) UMAP plot, color-coded by embryonic day (red: E8.0, light blue: E8.25). (I) UMAP plot showing pseudotime calculated from RNA velocity analysis.(J) PAGA graph illustrating the predicted differentiation trajectory from CPM to endothelial cells based on RNA velocity.

**Supplemental Figure 6**

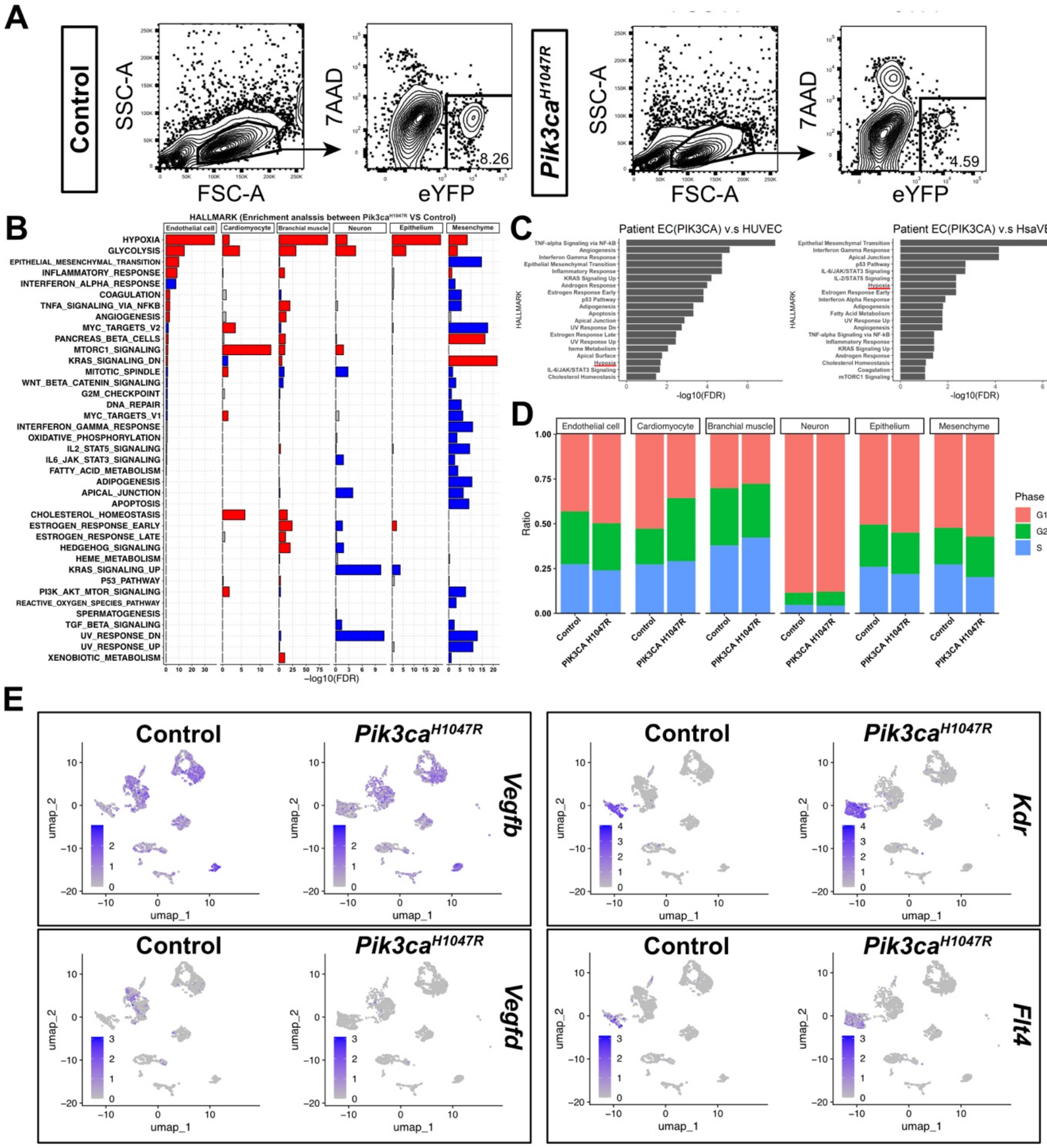

**Supplemental Figure 6. Pik3ca$^{\mathbf{H1047R}}$ expression enhances hypoxia signaling across multiple cell types**

**(A, B, D, E)** scRNA-seq analysis of eYFP$^{+}$ cells sorted by FACS from *Isl1-cre;R26R-eYFP* and *Isl1-cre;R26R-Pik3ca*$^{H1047R}$*;R26R-eYFP* embryos. **(A)** FACS sorting of eYFP+ cells. **(B)** Enrichment analysis from scRNA-seq comparing different cell types. Each bar chart shows Hallmark gene sets that exhibited significant changes in different cell types when comparing control and mutant groups. Red bars indicate Hallmark gene sets with higher enrichment scores in the mutant group, blue bars indicate higher scores in the control group, and gray bars indicate non-significant gene sets. The x-axis represents -log10(FDR), and significant Hallmark gene sets were defined as FDR < 0.1. **(C)** Re-analysis of bulk-RNA-seq data[25] from endothelial cells derived from PIK3CA mutated venous malformations (VM) (Patient EC(PIK3CA)) compared to control endothelial cells (Human umbilical venous endothelial cells: HUVEC: or Human saphenous vein endothelial cells: HsaVEC). The left bar chart shows the top 20 significantly enriched Hallmark gene sets in *PIK3CA* mutated Patient ECs compared to HUVEC, while the right chart shows enrichment in *PIK3CA* mutated Patient ECs compared to HsaVEC. Differentially expressed genes were defined as having a fold change >1.5 and FDR < 0.05. The x-axis represents -log10(FDR), and significant Hallmark gene sets were defined as FDR < 0.1. **(D)** Proportions of cells in each phase of the cell cycle (G1/S/G2-M) across cell types. **(E)** UMAP plots showing expression levels of *Vegfb, Vegfd, Kdr*, and *Flt4* by condition.

**Supplemental Figure 7**

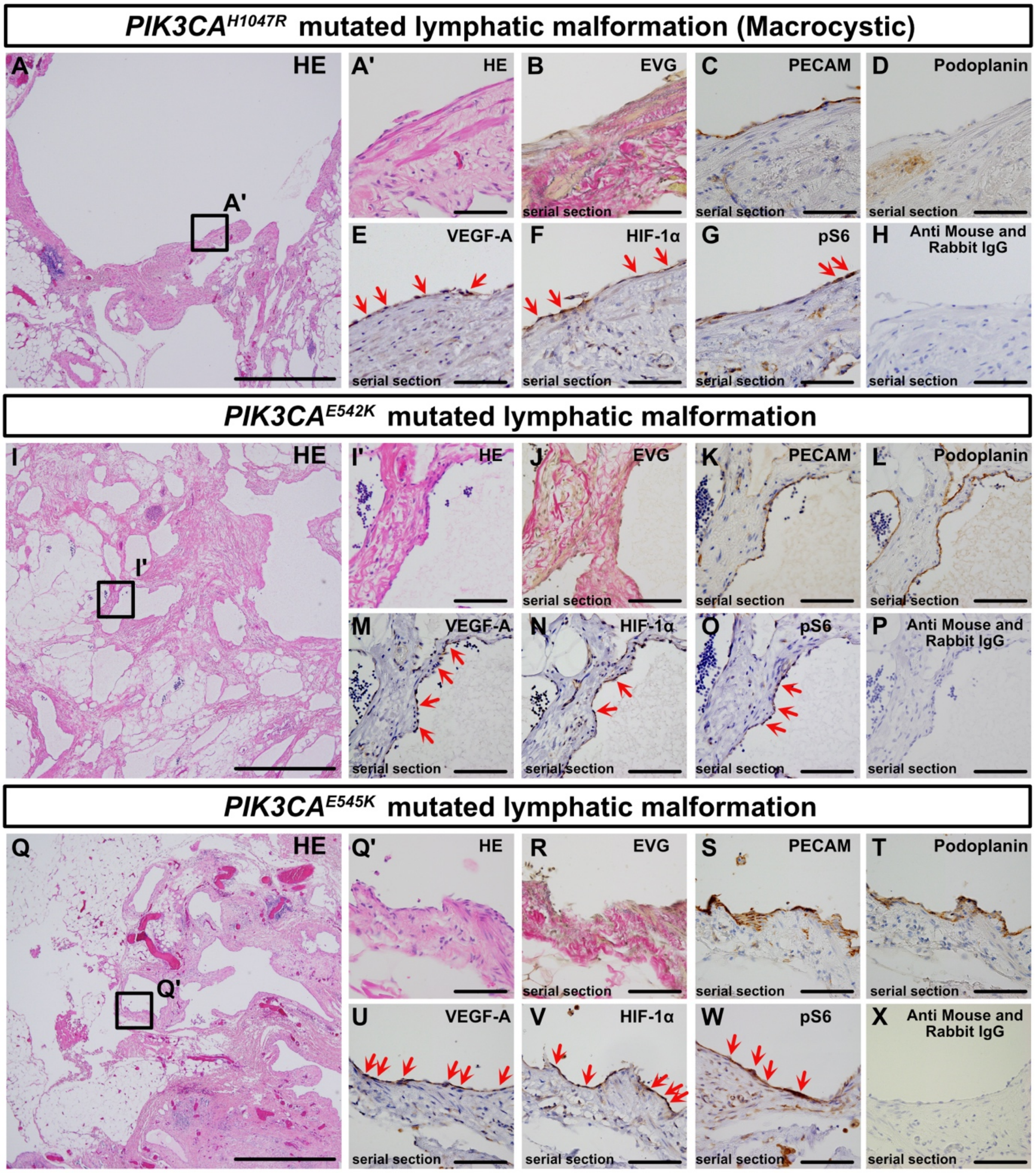

**Supplemental Figure 7. HIF-1α and VEGF-A expression in malformed lymphatic endothelial cells with *PIK3CA*$^{E542K}$ and *PIK3CA*$^{E545K}$ mutations.**

(**A-X**) Hematoxylin and eosin (HE) staining, special stains [elastica van Gieson (EVG), elastica Sirius red (ESR)—collagen fibers appear red, elastic fibers black, muscle tissue yellow], and immunohistochemistry using the indicated antibodies. No signal was detected in the negative controls using only secondary antibodies (**H, P, X**). (**A-H**) Macrocystic type lymphatic malformations. VEGF-A, HIF-1α, and pS6 are expressed in the ECs of malformed lymphatic vessels (**E-G**, red arrows). (**I-P”””**) Similar expression patterns are observed in lymphatic malformations with the *PIK3CA*$^{E542K}$ mutation, where VEGF-A, HIF-1α, and pS6 are detected in malformed LECs (**M-O**, red arrows). (**Q-X**) The same findings are present in lymphatic malformations with the *PIK3CA*$^{E545K}$ mutation, showing VEGF-A, HIF-1α, and pS6 expression in the malformed endothelial cells (red arrows, U-X). **Scale bars**: 100 μm (**A’-H, I’-P, Q’-X**), 1 mm (**A, I, Q**).
